# Early life stress leads to an aberrant spread of neuronal avalanches in the prefrontal-amygdala network in males but not females

**DOI:** 10.64898/2026.03.19.712827

**Authors:** Kharybina Zoia, Palva Matias, Palva Satu, Lauri Sari, Hartung Henrike, Taira Tomi

## Abstract

Development of the brain networks is highly vulnerable to stressful events. Early life stress (ELS) has been linked to multifaceted cognitive and emotional deficits in adulthood. Despite a growing body of evidence showing ELS-induced structural and functional changes in the prefrontal cortex (PFC) and basolateral amygdala (BLA), a circuit crucial for emotional processing, our knowledge of the resulting changes in the network dynamics is incomplete. Here, we investigate how maternal separation (MS) affects prefrontal-amygdala network in terms of neuronal avalanches, spatiotemporal clusters of activity, using simultaneous multielectrode recordings in the medial PFC (mPFC) and the BLA of urethane-anaesthetized juvenile (postnatal day (p) 14 – p15) and young adult (p50 – p 60) rats. Firstly, we show that MS leads to an intensified spread of activity within both regions as reflected in the higher mean branching ratios of the avalanches. Next, we demonstrate that most of the avalanches occur locally in one region, however, a small percentage of avalanches has clusters of activity in both regions simultaneously. We show that in MS animals prefrontal clusters followed by activity in the amygdala tend to be larger compared to controls and each event in the mPFC is followed by smaller number of events in the BLA, pointing towards impaired spread of activity from the mPFC to the BLA. Interestingly, avalanche spread from the BLA to the mPFC remains unaffected by MS. Abovementioned effects manifest only in adulthood and, intriguingly, only in males highlighting prolonged developmental and sex-dependent nature of ELS outcome.

**Significance statement:** Brain criticality implies that the brain self-organizers towards critical state, characterized by sustained activity propagation reflected in the unitary branching ratios of neuronal avalanches. Here we show how adverse events during early periods of network maturation, namely ELS, can disrupt developmental trajectories of the critical dynamics in the mPFC-BLA circuit in a sex-specific manner. This study broadens our understanding of the critical dynamics emergence in the prefrontal-limbic network and highlights ELS as a potential criticality control parameter.

## Introduction

Early periods of life are characterized by intense development and experience dependent refinement of neuronal networks in the brain (Tau and Peterson 2010; Dennis and Thompson 2013; De Benedictis et al. 2023), processes highly vulnerable to adverse events. Early life stress (ELS) has been linked to emotional, cognitive and reward processing deficits in adulthood (reviewed in Lupien et al. 2009; Pechtel and Pizzagalli 2011). People with history of ELS exposure have higher risk for depression (Pirkola et al. 2005), post-traumatic stress disorder (Famularo et al. 1992) and neurodevelopment disorders, such as attention deficit hyperactivity disorder and autism spectrum disorder (reviewed in Makris et al. 2023).

ELS has a profound effect on brain regions with protracted development, such as hippocampus, amygdala, corpus callosum, cerebellum, cortex (Pechtel and Pizzagalli 2011). Particularly vulnerable to ELS are amygdala and prefrontal cortex (PFC), regions rich in stress hormone receptors and playing a crucial role in emotion regulation (Tottenham 2020).

Early life adversity has been linked to the amygdala hyperactivity (McCrory et al. 2011; Tottenham et al. 2011; McCrory et al. 2012; Cohen et al. 2013; Gee et al. 2013b; Van Harmelen et al. 2013; Ishikawa et al. 2015; Yamamoto et al. 2017), increased amygdala volume (Mehta et al. 2009; Tottenham et al. 2010), but see (Edmiston et al. 2011; Hanson et al. 2015), and precocious myelination of axons in the basolateral amygdala (BLA) (Ono et al. 2008). In addition, chronic stress was shown to increase dendritic arborization in BLA (Vyas et al. 2002; Mitra et al. 2005). Concerning PFC, ELS was shown to have mostly opposite effects: reduced firing activity (Donati et al. 2026), hypoactivation (Levine et al. 2007; Levine et al. 2008; Van Harmelen et al. 2014; Ishikawa et al. 2015), decreased myelination (Makinodan et al. 2012; Yang et al. 2017), reduced dendritic length and branching (Pascual and Zamora-León 2007; Chocyk et al. 2013), but see (Muhammad et al. 2012), and smaller PFC volumes (Van Harmelen et al. 2010; Edmiston et al. 2011; Hanson et al. 2012; McLaughlin et al. 2014). Amygdala-prefrontal interactions were also shown to be affected by ELS which leads to accelerated maturation of functional (Gee et al. 2013b; Posner et al. 2016; Haikonen et al. 2022; Donati et al. 2026) and structural (Honeycutt et al. 2020; Nieves et al. 2020; Haikonen et al. 2022; Song et al. 2025) connectivity.

Given the above-mentioned structural and functional changes in BLA and PFC, ELS should have a great impact on the dynamical state of these regions. However, ELS-induced changes in the network dynamics of amygdala-prefrontal circuit remain poorly understood. Here, we address this issue in terms of the theory of self-organized criticality (Bak et al. 1987). Brain criticality implies that a healthy brain evolves towards critical point (Cocchi et al. 2017; Heiney et al. 2021), which is characterized by a stable propagation of activity through the system. Deviations from criticality disrupt efficient information transfer and are associated with cognitive deficits and mental diseases (Cocchi et al. 2017).

Previously (Kharybina et al. 2026) we described changes in the dynamical states of the medial PFC (mPFC) and BLA under normal development utilizing neuronal avalanches approach introduced by Beggs and Plenz (Beggs and Plenz 2003) and widely used to assess criticality in the brain. Current study investigates how ELS affects neuronal avalanches in the developing mPFC and BLA. We exploit maternal separation (MS), a well-established ELS model (Rincel and Darnaudéry 2020; Zhang et al. 2024) which implies daily separation of offsprings from their dam in the early postnatal period, and assess its effect on the mPFC-BLA network dynamics in rats in vivo at two developmental stages, juvenescence and young adulthood.

## Materials and methods

### Animals

All the procedures were carried out in accordance with the University of Helsinki Animal Welfare Guidelines and were approved by ELLA, the National Animal Experiment Board in Finland. Experiments were conducted on Han-Wistar rats, both sexes. Group sizes were determined based on previous experience.

Animals were in the temperature and humidity-controlled environment, with ad libitum access to food and water and 12/12h light/dark cycle. Pups from the litters were randomly assigned to two experimental groups with equal sex distribution. Pups from maternal separation (MS) group were separated from their litter and the dam for 3 hours (10 a.m – 1 p.m) per day from p2 to p14. During the separation, they were kept in individual chambers with heat pads to maintain body temperature. Littermate controls remained in their home cage with the dam.

Animals from both MS and control groups were divided into juvenile and adolescent groups. Experiments on the juvenile group were held on p14 – p15, the animals were sacrificed after the experiments. Pups from the adolescent group were weaned on p21 and then group housed (2-3 animals of the same sex in a cage). Experiments on the adolescent group were held on p50 – p60. We did not control for the Estrous cycle stage in females.

In total we recorded from 171 animals (19 litters). 8 animals were excluded from the analyses: 4 animals died during the recordings and 4 had low quality of the signal (artifacts or large noise level).

### In vivo electrophysiology

Detailed description of the surgery, electrophysiology recordings and histology is provided in (Kharybina et al. 2026).

In short, recordings were done under urethane anaesthesia (10 % urethane diluted in NaCl, i.p. injection 13 µl/g, top up 4 µl/g whenever needed). We used 4 % isoflurane for anaesthesia induction and 1 – 2 % isoflurane continuously for surgically deep anaesthesia. Two small burr holes were drilled above the regions of interest (mPFC and BLA) and one above the cerebellum. The body temperature was kept at 37°C during surgery and recordings.

For recordings, we used 32 (four-shank 4x8) or 64 (eight-shank 8x8) channel silicon multielecrode probes (NeuroNexus Technologies), 200 µm x 200 µm distance between the channels. Animals were fixed into stereotaxic frame and electrodes were inserted into the mPFC (four-shank probe, shanks parallel to the midline and perpendicular to the skull, 0.3 mm from the midline, 0.5 – 1.1 mm and 1.5 – 2.1 mm anterior to the bregma for the juvenile and adolescent animals, respectively, 3.5 mm depth) and BLA (four-shank probe for the juvenile and part of the adolescent animals, eight-shank probe for most of the adolescent animals, shanks were inserted at the base of the rhinal fissure at 40° from the vertical plane and perpendicular to the midline, 4.3 – 4.7 mm and 5 – 5.5 mm depth for the juvenile and adolescent animals, respectively). A reference silver wire was inserted into the cerebellum.

After at least 15 min recovery we performed extracellular recordings for 45 minutes with 20 kHz (Digital Lynx 4SX, Neuralynx) and 32 kHz sampling rate (Open Ephys acquisition board, Open Ephys).

### Data analysis

Only channels confined in the BLA and prelimbic part (PL) of the mPFC (deep and superficial layers pulled together) as confirmed by postmortem histological examination were used for further analyses. Data was analyzed with custom-written MATLAB (R2020b, Mathworks) scripts.

First, signal traces were preprocessed (third-order butterworth low-pass filter, <1000 Hz, downsampling to 2 kHz, IIR notch-filter at 50 Hz, q-factor = 35).

Based on histology and signal-to-noise ratio, 16 nearby channels in the regions of interest were picked for avalanche analysis. Avalanches were defined as clusters of negative deflections of LFP (nLFPs). To obtain LFP time courses, signals were normalized and band-pass filtered (third-order Butterworth filter, 1 – 100 Hz for the BLA traces and 2 – 100 Hz for the mPFC traces because of the prominent anaesthesia-induced high amplitude slow-rhythm in the mPFC). Then we detected nLFPs as maximum negative deflections of LFP after crossing a threshold and before crossing it back. We used a set of thresholds from 1 *sd* to 4 *sd* with the step of 0.2 *sd*, where *sd* is a standard deviation of the individual channels. Time scale was divided into the bins (time bin duration was in the range 1 – 30 ms with the step 0.2 ms). Subsequent time bins with nLFPs comprised one neuronal avalanche. Avalanches were separated from each other by at least one empty bin. Avalanches were characterized by the size s (a number of nLFPs in the avalanche) and branching ratio σ (ratio of nLFPs numbers in the second and the first bin of the avalanche). Experiments with less than 16 channels in the region of interest were discarded from the avalanche analysis. In total, mPFC and BLA avalanches were calculated for 123 and 110 animals, respectively. Avalanches spread between the regions was assessed only for one set of the parameters (2 *sd* threshold and 20 ms time bin) and in the experiments with at least 16 channels in both mPFC and BLA (in total, 86 animals).

Power and phase locking value were calculated using a continuous wavelet transform (complex Morlet transform, 5 cycles length, 50 logarithmically distributed center frequencies in the band 0.1 – 100 Hz). Calculations were done for the same 16 channels used in the avalanche analysis and results were averaged across these channels (power) or across all pairs of these channels (PLV).

For the detrended fluctuation analysis (DFA) we picked 5 nearby channels in the regions of interest based on histology and signal-to-noise ratio. For each channel we performed wavelet transform (complex Morlet wavelets, 5 cycles length, 20 logarithmically distributed center frequencies in the band 1 – 100 Hz). Then fluctuation function was calculated as described in (Hardstone et al. 2012) with window sizes logarithmically distributed from 1 s to 500 s and 80 % overlap. To get DFA exponents we calculated a slope of fluctuation function in double-logarithmic scale (least-squares method, linear fit from 5 s to 300 s). Results were averaged over the channels. Experiments with less than 5 channels in the region of interest were discarded from the analysis. In total, DFA exponents in the mPFC and BLA were calculated for 144 and 145 animals, respectively.

For more detailed description of the methods see (Kharybina et al. 2026).

For statistics, we ran separate Wilcoxon rank-sum test for all parameters (either frequency for power, PLV and DFA analyses or pair of threshold and bin size for avalanche analysis) and controlled for false discovery rate using Storey FDR correction which provides higher power than Benjamini-Hochberg method (Storey 2002). If not stated otherwise, we accepted q-values less than 0.05. To estimate an effect size, we calculated a probabilistic index as described in (Acion et al. 2006) and reported a maximum probabilistic index for all significant values clustered together.

To compare avalanche rates for avalanches with different localizations we used two-way repeated measures ANOVA with localization as repeated factor and age as a second factor. To compare ratios of small and large size probability and ratios between sizes of BLA and mPFC nLFP clusters we used two-way ANOVA with age and stress as factors. To assess stress effect on avalanche rate we used two-way ANOVA with stress and sex as factors. In all cases we used Holm-Sidak all pairwise comparison.

The sample size was based on our previous in vivo MS study (Haikonen et al. 2022).

### Data and code availability

The original data and original data analysis tools will be shared upon request.

## Results

### Effects of early life stress on network dynamics in the mPFC

Here we investigated how early life stress in the form of maternal separation (MS) affects mPFC network dynamics at different developmental stages. For this, we performed local field potential (LFP) recordings in urethane anaesthetized juvenile (postnatal day (p) 14 – p15) and young adult (p50 – p60) rats. Pups in MS group were daily separated from the dam and littermates for three hours at p2 – p14. As a control group we used their littermates kept all the time with the dam. Behavioral validation of the model was done previously in (Haikonen et al. 2022). In short, MS-exposed adult males spent less time in the central zone of the arena in the open field test, and in the open arms in the elevated plus maze test, consistent with anxiety-like behavioral phenotype. Travel distance in the open field test was reduced for MS males compared to the controls. No behavioral differences were found between MS and control females.

First, we checked whether MS had an effect on signal characteristics in the mPFC. Typical for sleep-like activity under urethane anaesthesia (Clement et al. 2008), in both control and MS groups in juvenile animals power spectra were characterized by peaks around 0.3 – 0.4 Hz, 2 – 4 Hz and 20 – 40 Hz (Figure S1A, S1D), in young adults there was a sharp peak around 1 Hz (Figure S1G, S1J). A similar developmental shift in peak delta frequency was reported in urethane anaesthetized mice (Donati et al. 2026). MS did not have significant effect on power spectra in males. In young adult females in MS group there was an increase in the power for infra slow rhythm 0.1 – 0.35 Hz (p < 0.05, Wilcoxon rank sum test, false discovery rate (FDR) correction, q < 0.2, N = 15 and 19 for control and MS groups, respectively). Note, however, that these results were accepted with 20% of false positives and need to be taken with some caution.

Next, we assessed functional connectivity within the region by calculating phase locking value (PLV) for all pairs of channels (Figure1, S1B, S1E, S1H, S1K). Close distance between recording sites (200 µm) and signal resemblance ensured high PLV values up to 0.9. These values were not affected by MS.

Our next question was whether MS affected long-range temporal correlations (LRTCs) in the mPFC, or the capacity of the system to remember its previous state. For this, we performed detrended fluctuation analysis (DFA) as described in (Hardstone et al. 2012). Resulting DFA exponents can range from 0 to 1 with values less than 0.5 reflecting negative LRTCs and values higher than 0.5 – positive LRTCs. In both MS and control groups, DFA exponents in juvenile animals were close to 0.5 as seen in the signals without LRTCs and went higher in young adults (Figure S1C, S1F, S1I, S1L). In other words, in both control and MS animals positive LRTCs in the mPFC emerged only in early adulthood. In both juvenile and young adult animals, DFA exponents did not differ between control and MS groups.

Then, we looked at the spatio-temporal characteristics of neuronal activity. For this, we calculated negative deflections of local field potentials (nLFPs) on 16 nearby channels. In both control and MS groups, nLFPs formed spatio-temporal clusters known as neuronal avalanches (Beggs and Plenz 2003). We detected avalanches for a set of parameters with the threshold of nLFPs detection in the range 1 – 4 standard deviations (SD) of the signal with the step of 0.2 SD and time bin in the range 1 – 30 ms with the step of 0.2 ms. For each avalanche we calculated its size as a number of nLFPs comprising an avalanche. Figure 1A, 1D depicts avalanche size distributions for males in the example case of 2 SD threshold and 2 ms time bin. Distributions of avalanche sizes followed truncated power-law with cut-off at 16 channels corresponding to the maximum number of channels used for avalanche detection and particularly prominent for small bin sizes. Distributions calculated with larger sizes were characterized by heavier tails and small bumps at 16 channels. This behavior was similar for both control and MS groups in both juvenile and adult animals.

**Figure 1.**
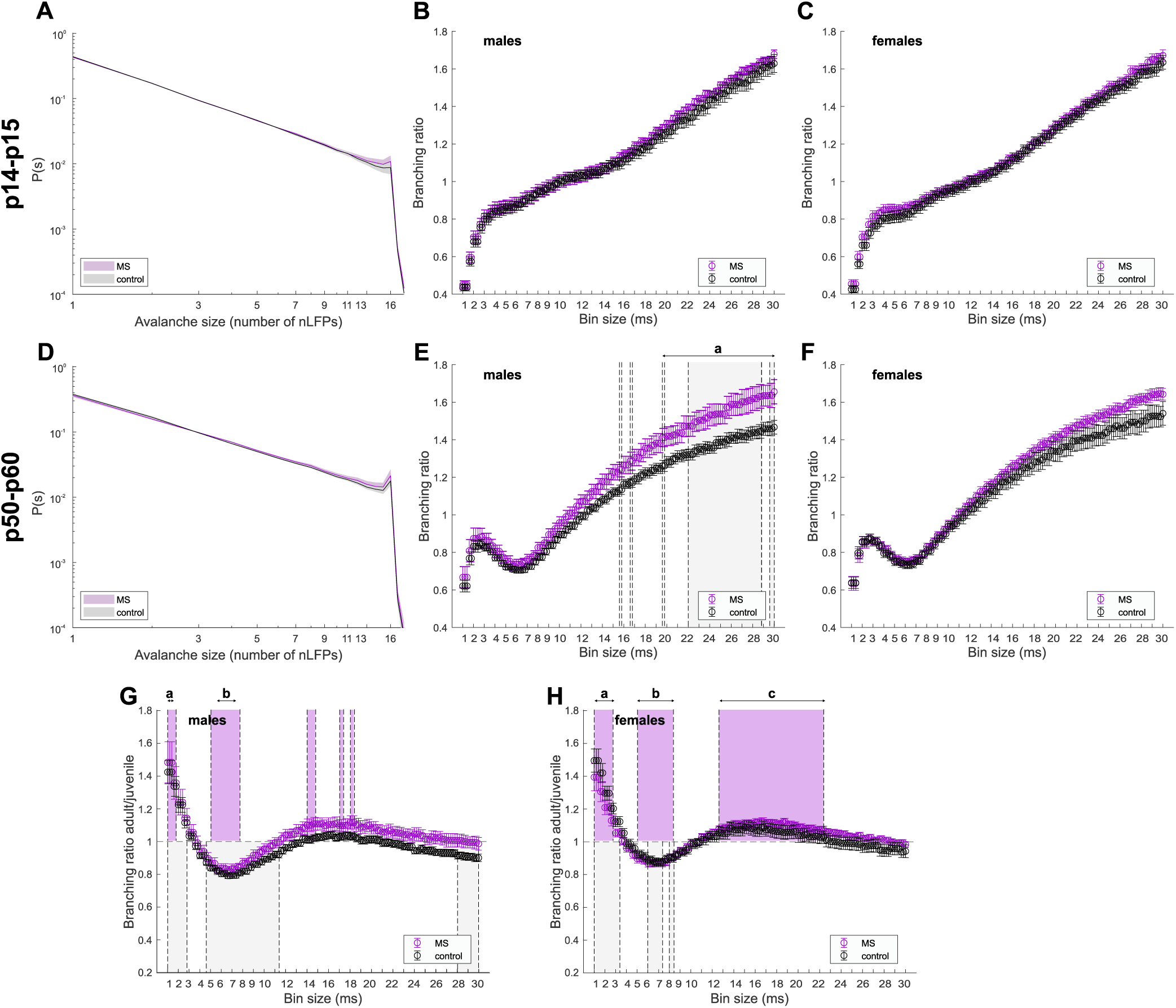
Effect of early life stress on neuronal avalanches in the mPFC. (A-C) Results for juvenile (p14 - p15) animals from MS (*magenta*) and control (*black*) groups. Shaded area (A) and error bars (B,C) represent SEM. In males and females N = 15 for both control and MS groups. (A) Avalanche size distribution calculated with 2SD threshold and 2 ms time bin averaged across males. For both control and MS groups size distributions follow truncated power-law with cut-off at 16 channels. (B) Branching ratio calculated with 2SD threshold and multiple values of time bin, averaged across males. (C) Same as (B) for females. (D-F) Same as (A-C) for young adult (p50 - p60) animals. In males: N = 17, 12; in females: N = 15, 19 for control and MS group, respectfully. In adult animals MS leads to significant increase in branching ratio for large bin sizes (grey rectangles in (E), p < 0.05, Wilcoxon rank sum test with FDR correction, q < 0.2) in males (E) but not females (F). (G, H) Branching ratio of adult animals calculated with 2SD threshold and multiple values of time bin, averaged across control (*black*) and MS (*magenta*) males (G) and females (H) and divided by mean branching ratio of juvenile animals from respective group. Grey and magenta rectangles show significant developmental differences (p < 0.05, Wilcoxon rank sum test with FDR correction, q < 0.05) for control and MS groups, respectively. In both groups and sexes branching ratio increases with age for small bin sizes less than 3 ms and decreases for bin sizes between 5 and 9 ms. In males, the effect is less pronounced in MS group. In females, in MS group there is an additional developmental increase in the branching ratio for large bin sizes between 12 and 23 ms.

Next, we calculated a branching ratio, i.e. a ratio of the numbers of nLFPS in the second and the first bins of an avalanche, and averaged it across all avalanches in the region. Mean branching ratio shows how many events are expected to be triggered by a single event. For the systems in the critical state this parameter is close to 1. In general, mean branching ratio dependance on the threshold and bin size in MS animals closely resembled that for control animals, monotonically increasing with larger bin size and decreasing with larger threshold in juveniles and having additional peak at bin size around 3 ms in young adult animals (Figure S2A, S2C). In juvenile animals we failed to see a difference in the mean branching ratio between control and MS groups (Figure 1B, 1C, S2B, S2D). In young adult males mean branching ratio was significantly higher (p < 0.05, Wilcoxon rank sum test, FDR correction, q < 0.2, N = 17 and 12 for control and MS groups, respectively) for MS animals for a large set of parameters with bin size higher than 10 ms. In Figure 1E, S2D significant differences in the branching ratio are shown after FDR correction with q < 0.2. Most of the points in the parameter space with significant difference formed one large cluster (Figure S2D, cluster **a**, size 624). To characterize an effect size, for each point in the cluster we calculated a probability index P(MS>control) showing the probability of the mean branching ratio in the MS group being higher than that in the control group. Maximum value across all the points in the cluster **a** *maxP(MS>control)* was 0.77. In young adult females we saw the same trend (Figure 1F, S2D), however difference between the groups did not reach significant levels.

Then we looked at the developmental changes in the avalanche dynamics under MS. In males, for both control and MS groups we saw the same developmental changes (Figure 1G, S2E): an increase in the branching ratio for small bin sizes (cluster **a**: cluster size 152 and 125, *maxP(adult>juvenile)* = 0.9 and 0.89 for control and MS group, respectively; p < 0.05, Wilcoxon rank sum test, FDR correction, q < 0.05, N = 15 and 17, 15 and 12, for control juvenile and young adult males, and MS juvenile and young adult males, respectively) and decrease for bin sizes between 5 and 11 ms (cluster **b**: cluster size 372 and 170, *maxP(adult<juvenile)* = 0.95 and 0.97 for control and MS group, respectively), however, with smaller cluster sizes in MS group. In other words, developmental changes in the avalanche characteristics in the MS males were less pronounced compared to the normal development. Interestingly, while we failed to see MS effect in females in any age group, developmental changes in the MS females were more striking than in the controls. Whereas in the control females we saw only a significant increase for small bin sizes (cluster **a**: cluster size 208, *maxP(adult>juvenile)* = 0.96; p < 0.05, Wilcoxon rank sum test, FDR correction, q < 0.05, N = 15 for both juvenile and young adult females) and decrease for larger bins was not significant, in the MS females both these clusters were significant (cluster **a**: cluster size 158, *maxP(adult>juvenile)* = 0.88; cluster **b**: cluster size 154, *maxP(adult<juvenile)* = 0.89; p < 0.05, Wilcoxon rank sum test, FDR correction, q < 0.05, N = 15 and 19 for juvenile and young adult females, respectively) (Figure 1H, S2E). Moreover, in MS females there was a significant developmental increase in the branching ratio for a large cluster of parameters (cluster c: cluster size 602, *maxP(adult>juvenile)* = 0.87) with bin sizes between 12 and 25 ms. A similar increase for large bin sizes was seen in control females and MS males, but it was not significant.

To sum up, in the mPFC in males in both age groups MS had no effect on LFP power, however there was an increase in young adult MS females in infra-slow band compared to the control females. MS did not affect inter-regional functional connectivity and LRTCs in either age group. Neuronal avalanches in juvenile animals were not affected by MS. However, MS led to the significant increase in the mean branching ratio of prefrontal neuronal avalanches detected with large bin sizes in young adult males but not females. Developmental changes in the avalanche branching ratio in the mPFC were less pronounced in MS males compared to the normal development, whereas in MS females these changes were more distinct.

### Effects of early life stress on network dynamics in the BLA

Here we investigated how MS affects network dynamics in the BLA. First, we analyzed signal characteristics in the BLA. Same as in the mPFC, mean power spectra calculated for 16 adjacent channels in BLA in juvenile animals were characterized by peaks around 0.3 – 0.4 Hz, 2 – 4 Hz and 20 – 40 Hz (Figure S3A, S3D). In young adult animals, there was a peak around 1 Hz (Figure S3G, S3J) but less sharp than in the mPFC. We failed to see significant effects of MS on power spectra in both age groups. Inter-regional functional connectivity in the BLA assessed by PLV was not affected by MS as well (Figure S3B, S3E, S3H, S3K).

We proceeded by characterizing LRTCs in BLA. Both in control and MS animals LRTCs in the BLA emerged already in juvenile animals (Figure S3C, S3F) with DFA exponents being above 0.5 for frequencies up to 60 Hz and going even higher in young adult animals (Figure S3I, S3L). MS did not have a significant effect on DFA exponents in both age groups.

The next question was the effect of MS on neuronal avalanches in the BLA. Both in control and MS group avalanche size distributions in the BLA did not follow power law and were instead characterized by a sharp peak at 16 channels with larger peak in case of young adult animals (Figure 2A, 2D).

**Figure 2.**
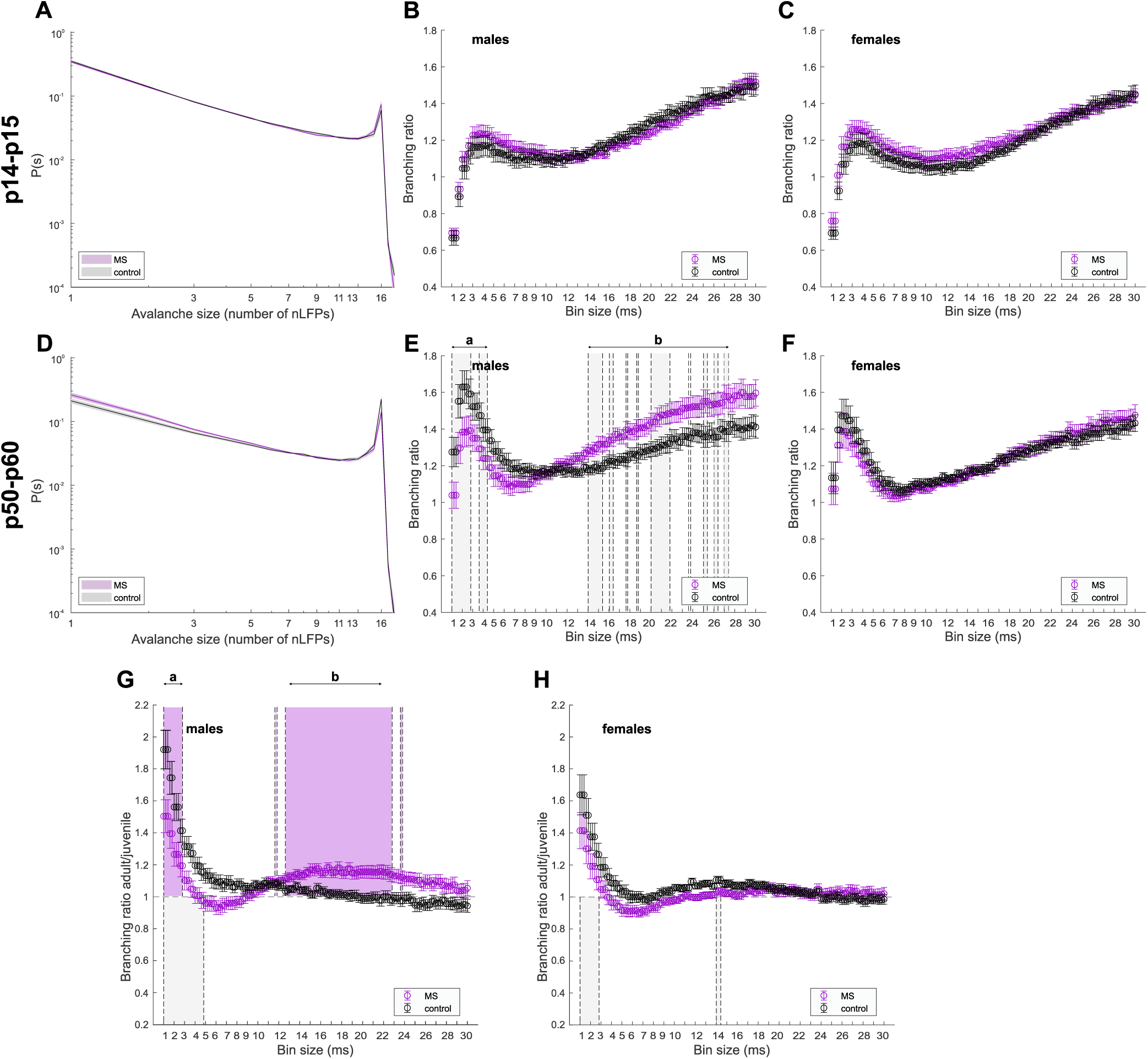
Effect of early life stress on neuronal avalanches in the BLA. (A-C) Results for juvenile (p14 - p15) animals from MS (*magenta*) and control (*black*) groups. Shaded area (A) and error bars (B,C) represent SEM. In males: N = 11, 13; in females: N = 17, 11 for control and MS group, respectfully. A. Avalanche size distribution calculated with 2SD threshold and 2 ms time bin averaged across males. For both control and MS groups size distributions deviate from power-law and have sharp peak corresponding to 16 channels. (B) Branching ratio calculated with 2SD threshold and multiple values of time bin, averaged across males. (C) Same as (B) for females. (D-F) Same as (A-C) for young adult (p50 - p60) animals. In males: N = 13, 12; in females: N = 15, 18 for control and MS group, respectfully. In adult animals MS leads to significant decrease in branching ratio for small bin sizes and increase for large bin sizes (grey rectangles in (E), p < 0.05, Wilcoxon rank sum test with FDR correction, q < 0.05) in males (E) but not females (F). (G, H) Branching ratio of adult animals calculated with 2SD threshold and multiple values of time bin, averaged across control (*black*) and MS (*magenta*) males (G) and females (H) and divided by mean branching ratio of juvenile animals from respective group. Grey and magenta rectangles show significant differences (p < 0.05, Wilcoxon rank sum test with FDR correction, q < 0.05) for control and MS groups, respectively. There is a developmental increase in the branching ratio for small bin sizes less than 5 ms, less pronounced in MS males compared to controls and insignificant in MS females. In MS group there is an additional increase in the branching ratio for large bin sizes between 13 and 22 ms in males (G) but nor females (H).

In the BLA, mean branching ratio dependance on the threshold and bin size is similar in control and MS animals. It is larger than one for most of the threshold values, has a peak around 3-4 ms and monotonically increases for larger bin sizes (Figure 2B, 2C, 2E, 2F, S4A, S4C). In juvenile animals, both males and females, MS did not affect mean branching ratio (Figure 2B, 2C, S4B), whereas in young adult males we saw two distinct clusters of parameters with significant differences in the mean branching ratio (Figure 2E, S4D) (p < 0.05, Wilcoxon rank sum test with FDR correction, q < 0.05, N = 13 and 12 for control and MS groups, respectively). For small bin sizes less than 5 ms (cluster **a**: size 245, maxP(MS<control) = 0.83) MS resulted in the decreased branching ratio. However, for bin sizes larger than 13 ms (cluster **b**: size 953, *maxP(MS>control)* = 0.86) the branching ratio was higher in MS compared to the controls. In young adult females we saw the same trend, but differences were not significant (Figure 2F, S4D).

Next, we looked whether MS affected developmental changes in the avalanche dynamics in the BLA. Under control conditions there was a developmental increase in the branching ratio (p < 0.05, Wilcoxon rank sum test with FDR correction, q < 0.05) for small bin sizes (less than 5 ms), more prominent in males (cluster **a**: size 282, *maxP(adult>juvenile)* = 0.97; N = 11 and 13 for juvenile and young adult animals, respectively) than in females (cluster **a**: size 149, *maxP(adult>juvenile)* = 0.86; N = 17 and 15 for juvenile and young adult animals, respectively). In case of MS, we saw the same increase in males, though for smaller cluster of parameters (cluster **a**: size 148, *maxP(adult>juvenile)* = 0.88) limited by bin sizes less than 3 ms (Figure 2G, S4E) (p < 0.05, Wilcoxon rank sum test with FDR correction, q < 0.05, N = 13 and 12 for juvenile and young adult animals, respectively). In contrast to the control animals, in MS males we saw an additional large parameter cluster with significant increase in the branching ratio (cluster **b**: size 483, *<P(adult>juvenile)>* = 0.98) with bin sizes between 12 and 24 ms. In MS females we saw a developmental increase only for small bin sizes (Figure 2H, S4E), however this cluster did not pass FDR correction.

Overall, MS had no effect on power, inter-regional functional connectivity and LRTCs in the BLA. However, we saw MS driven changes in the characteristics of neuronal avalanches in the BLA in young adult animals. Interestingly, the effect valence depended on the bin size with MS leading to decrease in the mean branching ratio for small bin sizes and increase for large bin sizes. As a result, developmental increase in the branching ratio in case of small bin sizes was less pronounced in MS animals, however, we saw an additional increase in case of large bin sizes not seen in control animals. This effect was specific to male animals but not female.

### Effects of early life stress on avalanche spread between the mPFC and the BLA

Previously we have shown that a small part of avalanches spread between the mPFC and the BLA (Kharybina et al. 2026). Considering MS caused changes in the branching ratio in both mPFC and BLA we decided to check if MS affects avalanche spread between the regions.

In both control and MS animals, most of nLFPs were clustered locally either in the mPFC or in the BLA, however some of the nLFP clusters were detected in both regions simultaneously, forming, as we would call them, two-region avalanches (Figure 3A). For detailed description we focused on the avalanches calculated with 2SD threshold and 20 ms time bin, time comparable with the conductance delay between the mPFC and the BLA in adult rats (McGinty and Grace 2008), and calculated avalanche occurrence rate. Note, that nLFP clusters occurring in two regions simultaneously were counted as one avalanche.

**Figure 3.**
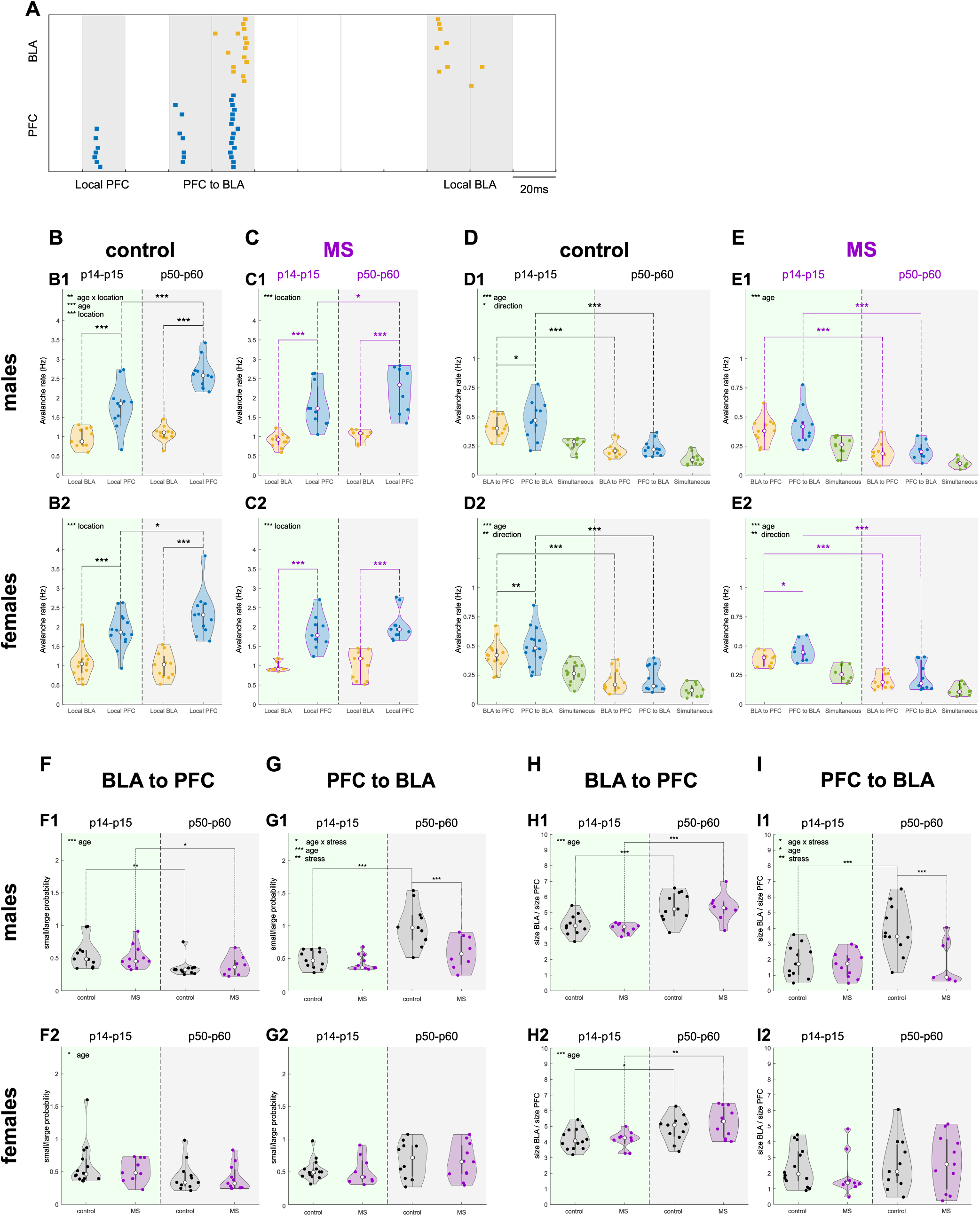
Spread of avalanches between the mPFC and the BLA. (A) Raster plot of nLFPs detected in BLA (yellow) and PFC (blue) of p14 male rat with 2SD threshold. Here nLFPs are clustered into three avalanches calculated with 20 ms time bin, two of which occur locally, either in the mPFC or the BLA, and one starts in the mPFC and then appears in the BLA (PFC to BLA avalanche). (B, C) Occurrence rate of local avalanches (2SD, 20 ms) for juvenile (green area) and young adult (grey area) control (B) and MS (C) males (B1, C1) and females (B2, C2). Main effects: there is significant age-location interaction in control males (B1: F = 9.343, P = 0.006), but not in other groups (B2: F = 3.308, P = 0.082, C1: F = 1.136, P = 0.301, C2: F = 0.219, P = 0.646); in control males main effects cannot be properly interpreted (B1: age F = 15.797, P < 0.001; location F = 103.265, P < 0.001), in control females and MS males occurrence rate of local avalanches is higher in PFC than in BLA (B2: F = 61.583, P < 0.001, C1: F = 61.380, P < 0.001, C2: F = 47.741, P < 0.001), there are no developmental changes in the occurrence rate (B2: F = 2.374, P = 0.137, C1: F = 3.446, P = 0.081, C2: F = 0.402, P = 0.534). All pairwise comparison: occurrence rate increases with development in PFC (B1: t = 4.971, p < 0.001, B2: t = 2.384, p = 0.021, C1: t = 2.090, p = 0.044) in all groups except MS females (C2: t = 0.745, p = 0.461), but not in BLA (B1: t = 0.642, p = 0.524, B2: t = 0.388, p = 0.7, C1: t = 0.653, p = 0.518, C2: t = 0.0231, p = 0.982); occurrence rate for local avalanches is higher in PFC than in BLA for both juvenile (B1: t = 5.024, p < 0.001, B2: t = 4.544, p < 0.001, C1: t = 5.216, p < 0.001, C2: t = 4.343, p < 0.001) and young adult animals (B1: t = 9.347, p < 0.001, B2: t = 6.459, p < 0.001, C1: t = 5.849, p < 0.001, C2: t = 5.499, p < 0.001). (D, E) Same as (B, C) for two-region avalanches. Main effects: there is no significant interaction between age and avalanche directionality (D1: F = 1.249, P = 0.277, D2: F = 0.966, P = 0.336, E1: F = 0.229, P = 0.638, E2: F = 0.633, P = 0.437); in all groups there is a developmental decrease in the occurrence rate of two-region avalanches (D1: F = 28.847, P < 0.001, D2: F = 28.612, P < 0.001, E1: F = 18.355, P < 0.001, E2: F = 25.222, P < 0.001) with higher avalanche rate for PFC to BLA avalanches in all groups (D1: F = 6.232, P = 0.021, D2: F = 12.535, P = 0.002, E2: F = 10.161, P = 0.005) except MS males (E1: F = 4.117, P = 0.058). All pairwise comparison: occurrence rate decreases with age for both PFC to BLA (D1: t = 5.465, p < 0.001, D2: t = 5.438, p < 0.001, E1: t = 3.997, p < 0.001, E2: t = 5.048, p < 0.001) and BLA to PFC (D1: t = 4.822, p < 0.001, D2: t = 5.060, p < 0.001, E1: t = 4.255, p < 0.001, E2: t = 4.612, p < 0.001) avalanches; occurrence rate is higher for PFC to BLA avalanches in juvenile animals for all groups (D1: t = 2.556, p = 0.019, D2: t = 3.410, p = 0.002, E2: t = 2.686, p = 0.015) except MS males (E1: t = 1.194, p = 0.249), but not in young adult animals (D1: t = 0.975, p = 0.341, D2: t = 1.709, p = 0.101, E1: t = 1.648, p = 0.118, E2: t = 1.783, p = 0.091). In (B-E) two-way repeated measures ANOVA on raw (B1, B2, C1) or log-transformed (D1, D2, C2, E1, E2) data with Holm-Sidak all-pairwise comparison. (F) Ratio between probability of small (s < 5) and large (17 < s < 50) sizes of BLA clusters in BLA to PFC avalanches in juvenile (green area) and young adult (grey area) control (*black*) and MS (*magenta*) males (F1) and females (F2). Main effects: there is no interaction between age and stress (F1: F = 0.424, P = 0.519, F2: F = 0.0294, P = 0.865); there is a developmental decrease in the small/large size probability ratio (F1: F = 13.560, P < 0.001, F2: F = 5.522, P = 0.024) with no effect of stress (F1: F = 0.00867, P = 0.926, F2: F = 0.279, P = 0.6). All pairwise comparison: there is a developmental decrease in the small/large size probability ratio in males for both control (F1: t = 3.205, p = 0.003) and MS (F1: t = 2.057, p = 0.047) groups and no developmental changes in females in both control (F2: t = 1.889, p = 0.066) and MS (F2: t = 1.463, p = 0.151) groups; no stress effect in both juvenile (F1: t = 0.550, p = 0.585, F2: t = 0.494, p = 0.624) and young adult (F1: t = 0.379, p = 0.707, F2: t = 0.253, p = 0.802) groups. (G) Same as (F) for PFC clusters in PFC to BLA avalanches. Main effects: there is an interaction between age and stress in males (G1: F = 4.346, P = 0.044) but not in females (G2: F = 0.00184, P = 0.966); in males main effects cannot be properly interpreted (G1: age F = 21.600, P < 0.001, stress F = 9.804, P = 0.003), in females there is no developmental (G2: F = 3.462, P = 0.07) nor stress effects (G2, F = 0.246, P = 0.966) in the small/large size probability ratio. All pairwise comparison: there is a developmental increase in the small/large size probability ratio in control (G1: t = 4.979, p < 0.001), but not MS (G1: t = 1.739, p = 0.09) males, and no developmental changes in both control (G2: t = 1.362, p = 0.181) and MS (G2: t = 1.278, p = 0.208) females; the small/large size probability ratio is decreased in males in MS animals in young adult (G1: t = 3.540, p < 0.001), but not juvenile (G1: t = 0.0477, p = 0.444) group, and unaffected in females in both juvenile (G2: t = 0.381, p = 0.705) and young adult (G2: t = 0.321, p = 0.75) groups. (H) Ratio between sizes of BLA clusters and PFC clusters from BLA to PFC avalanches in juvenile (green area) and young adult (grey area) control (*black*) and MS (*magenta*) males (H1) and females (H2). Main effects: there is no interaction between age and stress (H1: F = 0.206, P = 0.653, H2: F = 0.821, P = 0.370); there is a developmental increase in the BLA/PFC cluster size ratio (H1: F = 31.676, P < 0.001, H2: F = 14.835, P < 0.001) with no effect of stress (H1: F = 0.444, P = 0.509, H2: F = 0.565, P = 0.456). All pairwise comparison: there is a developmental increase in the BLA/PFC cluster size ratio for both control (H1: t = 3.826, p < 0.001, H2: t = 2.207, p = 0.033) and MS animals (H1: t = 4.127, p < 0.001, H2: t = 3.195, p = 0.003) and no stress effect in both juvenile (H1: t = 0.828, p = 0.413, H2: t = 0.109, p = 0.914) and young adult (H1: t = 0.144, p = 0.886, H2: t = 1.173, p = 0.247) animals. (I) Same as (H) for PFC to BLA avalanches. Main effects: there is a significant interaction between age and stress in males (I1: F = 6.157, P = 0.018) but not in females (I2: F = 0.231, P = 0.633); in males main effects cannot be properly interpreted (I1: age F = 6.792, P = 0.013; stress F = 8.551, P = 0.006), in females there is no developmental (I2: F = 0.178, P = 0.676) nor stress effects (I2, F = 0.782, P = 0.382) in BLA/PFC cluster size ratio. All pairwise comparison: there is a developmental increase in the BLA/PFC cluster size ratio in control (I1, t = 3.762, p < 0.001), but not MS (I1, t = 0.0848, p = 0.933) males, and no developmental changes in both control (I2: t = 0.0444, p = 0.965) and MS (I2: t = 0.606, p = 0.548) females; in males BLA/PFC cluster size ratio is decreased in MS animals in young adult (I1: t = 3.668, p < 0.001), but not juvenile (I1: t = 0.328, p = 0.745) group and unaffected in females in both juvenile (I2: t = 0.964, p = 0.341) and young adult (I2: t = 0.286, p = 0.777) groups. In (F-I) two-way ANOVA on raw (H2, I1) or log-transformed (F1, F2, G1, G2, H1, I2) data with Holm-Sidak all-pairwise comparison. In (B-I) in juvenile animals N = 11 in both control and MS males, N = 14, 9 in control and MS females, respectively; in young adult animals N = 11, 8 in control and MS males, respectively, N = 11 in both control and MS females.

First, we looked at the local avalanches. In juvenile animals there were no MS-related changes in the occurrence rate of local avalanches in both mPFC (main effect P = 0.896, two-way ANOVA) and BLA (main effect P = 0.35, two-way ANOVA) (Figure S5). In early adulthood, occurrence rate of local prefrontal avalanches was higher in MS animals (main effect P = 0.017, two-way ANOVA on log-transformed data). However, Holm-Sidak all-pairwise comparison showed no significant differences in neither males (p = 0.053) nor females (p = 0.140). We failed to see MS-related changes in the occurrence rate of local BLA avalanches in young adults (main effect P = 0.975, two-way ANOVA).

Then, we looked at the developmental changes in the occurrence of local avalanches (Figure 3B, 3C). In control males, there was significant interaction between age and location (age F = 15.797, P < 0.001; location F = 103.265, P < 0.001; interaction F = 9.343, P = 0.006, two-way repeated measures ANOVA), so main effects could not be properly interpreted, whereas in MS males there was no interaction between age and location (F = 1.136, P = 0.301), occurrence rate of local avalanches was significantly higher in the mPFC (F = 61.380, P < 0.001) with no developmental changes (F = 3.446, P = 0.081). Holm-Sidak all-pairwise comparison showed that in both control and MS males local avalanches in the mPFC occurred more often compared to the BLA in both age groups (controls: t = 5.024, p < 0.001 and t = 9.347, p < 0.001, MS: t = 5.216, p < 0.001 and t = 5.849, p < 0.001, p < 0.001 for juvenile and young adult males, respectively). With age, avalanche occurrence rate in the BLA remained the same (t = 0.642, p = 0.524 and t = 0.653, p = 0.518 for control and MS groups, respectively) whereas in the mPFC it increased in early adulthood with developmental changes in MS group being less pronounced (t = 4.971, p < 0.001 and t = 2.090, p = 0.044 for control and MS groups, respectively). Similar results were found in females, except that we failed to see significant developmental changes in the occurrence rate in the mPFC in MS group (t = 2.384, p = 0.021 and t = 0.745, p = 0.461 for control and MS groups, respectively).

Next, we looked at the two-region avalanches. First, we should clarify some definitions. Imagine that in one region we have an nLFP cluster A with its first nLFP in time bin t_1_ and its last nLFP in time bin t_end_. We would say that nLFP cluster B in the second region follows nLFP cluster A if the first nLFP of the cluster B is in any time bin from t_2_ to t_end+1_. Both nLFP clusters would be counted as one avalanche starting in the first region. For most of the two-region avalanches we could attribute directionality either from the mPFC to the BLA (PFC to BLA avalanches), meaning that they started from nLFPs in the mPFC and were followed by nLFPs in the BLA, or from the BLA to the mPFC (BLA to PFC avalanches). However, in some cases both nLFP clusters started in the same time bin (simultaneous avalanches).

In both control and MS animals, two-region avalanches constituted only a small proportion of the total number of avalanches. Comparing MS and control groups, we failed to see any MS-related changes in the occurrence rate of two-region avalanches of any directionality for both ages and sexes (Figure S5). Next, we looked at the developmental changes in the occurrence rate of two-region avalanches (Figure 3D, 3E). In males, the occurrence rate significantly decreased with development in both groups (F = 28.847, P < 0.001 and F = 18.355, P < 0.001 for control and MS males, respectively, two-way repeated measures ANOVA on log-transformed data, Holm-Sidak all-pairwise comparison) for both PFC to BLA (t = 5.465, p < 0.001 and t = 3.997, p < 0.001 for control and MS males, respectively) and BLA to PFC (t = 4.822, p < 0.001 and t = 4.255, p < 0.001 for control and MS males, respectively) avalanches. The same was true for developmental changes in both control and MS females. Results obtained for control animals showed a leading role of the mPFC in case of two-region avalanches (F = 6.232, P = 0.021 and: F = 12.535, P = 0.002 for control males and females, respectively), with PFC to BLA occurrence rate significantly higher compared with BLA to PFC avalanches in juveniles (t = 2.556, p = 0.019 and t = 3.410, p = 0.002, for control males and females, respectively) but not in young adults (t = 0.975, p = 0.341 and t = 1.709, p = 0.101, for control males and females, respectively). In MS males a leading role could not be attributed to any direction (main effect F = 4.117, P = 0.058; t = 1.194, p = 0.249 and t = 1.648, p = 0.118 for juvenile and young adult MS males, respectively). Interestingly, this loss of leading role was not seen in MS females (main effect F = 10.161, P = 0.005; t = 2.686, p = 0.015 and t = 1.783, p = 0.091 for juvenile and young adult MS females, respectively).

Next, we asked if there are any changes in the sizes of two-region avalanches. First, we computed size distributions of the starting nLFP clusters (Figure S6), i.e. prefrontal nLFP cluster sizes in PFC to BLA avalanches and amygdaloid nLFP cluster sizes in BLA to PFC avalanches, and noted that for PFC to BLA avalanches nLFP cluster sizes in the mPFC tended to have less small (s < 5) and more large (17 < s < 50) avalanches in young adult males in MS group. To estimate this, we calculated a ratio of probabilities of small and large cluster sizes (Figure 3F, 3G). In BLA to PFC avalanches we saw developmental decrease in this ratio (in males: main effect F = 13.560, P < 0.001; t = 3.205, p = 0.003 and t = 2.057, p = 0.47 for control and MS groups, respectively; in females: main effect F = 5.522, P = 0.024; t = 1.889, p = 0.066 and t = 1.463, p = 0.151 for control and MS groups, note, however, that results for each female group were not significant, two-way ANOVA on log-transformed data with Holm-Sidak all-pairwise comparison) but no MS-related differences (in males: main effect F = 0.00867, P = 0.926; t = 0.550, p = 0.585 and t = 0.379, p = 0.707 for juvenile and young adult groups, respectively; in females: main effect F = 0.279, P = 0.6; t = 0.494, p = 0.624 and t = 0.253, p = 0.802 for juvenile and young adult groups, respectively). Thus, LFP cluster transfer from the BLA to the mPFC was not affected by MS.

In PFC to BLA avalanches developmental changes in the probability ratio were found only in control males (in males: significant interaction between age and stress F = 4.346, P = 0.044, main age effect F = 21.600, P < 0.001; t = 4.979, p < 0.001 and t = 1.739, p = 0.09 for control and MS groups, respectively; in females: main age effect F = 3.462, P = 0.07; t = 1.362, p = 0.181 and t = 1.278, p = 0.208 for control and MS groups, respectively, two-way ANOVA on log-transformed data with Holm-Sidak all-pairwise comparison), where probability ratio increased with age. MS-related changes were observed only in young adult males (in males: main effect F = 9.804, P = 0.003; t = 0.0477, p = 0.444 and t = 3.540, p < 0.001 for juvenile and young adult groups, respectively; in females: main effect F = 0.246, P = 0.622; t = 0.381, p = 0.705 and t = 0.321, p = 0.75 for juvenile and young adult groups, respectively) with the probability ratio being significantly lower in MS animals. Thus, in MS animals for the nLFP clusters to spread from the mPFC to the BLA their size should be larger than in controls. The effect was specific for young adult males but not seen in females and juvenile animals.

Then we checked how many events in one region are triggered by one event in the other region. For this, for each two-region avalanche we calculated a ratio between sizes of nLFP clusters in the BLA and nLFP clusters in the mPFC (Figure 3H, 3I). In BLA to PFC avalanches this ratio is higher than one and increases with age (in males: main effect F = 31.676, P < 0.001; t = 3.826, p < 0.001 and t = 4.127, p < 0.001 for control and MS males, respectively; in females: main effect F = 14.835, P < 0.001; t = 2.207, p = 0.033 and t = 3.195, p = 0.003 for control and MS females, respectively, two-way ANOVA on log-transformed (males) and raw (females) data with Holm-Sidak all-pairwise comparison), meaning that on average a single event in the mPFC is preceded my several events in BLA, more so in young adult animals. No MS-related changes were found (in males: main effect F = 0.444, P = 0.509; t = 0.828, p = 0.413 and t = 0.144, p = 0.886 for juvenile and young adult males, respectively; in females: main effect F = 0.565, P = 0.456; t = 0.109, p = 0.914 and t = 1.173, p = 0.247 for juvenile and young adult females, respectively).

In PFC to BLA avalanches BLA/PFC nLFP cluster size ratio was also higher than one meaning that one event in the mPFC on average is followed by several events in the BLA. In females there were no developmental (main effect F = 0.178, P = 0.676; t = 0.0444, p = 0.965 and t = 0.606, p = 0.548 for control and MS groups, respectively, two-way ANOVA on log-transformed data with Holm-Sidak all-pairwise comparison) nor MS-related (main effect F = 0.782, P = 0.382; t = 0.964, p = 0.341 and t = 0.286, p = 0.777 for juvenile and young adult groups, respectively) changes in this ratio. In males, we saw developmental increase in this ratio in control but not MS group (significant interaction between age and stress F = 6.157, P = 0.018, main age effect F = 6.792, P = 0.013; t = 3.762, p < 0.001 and t = 0.0848, p = 0.933, for control and MS groups, respectively, two-way ANOVA with Holm-Sidak all-pairwise comparison). BLA/PFC nLFP cluster size ratio was lower in MS young adult, but not juvenile, males compared to controls (main stress effect F = 8.551, P = 0.006; t = 0.328, p = 0.745 and t = 3.668, p < 0.001 for juvenile and young adult groups, respectively).

Summing up, most of the avalanches occurred locally in one region. The occurrence of local PFC avalanches went up with age (but not in MS females), remained the same for local BLA avalanches and decreased for two-region avalanches (both PFC to BLA and BLA to PFC). In case of two-region avalanches in juvenile animals we could attribute preferred direction of avalanche spread with more PFC to BLA than BLA to PFC avalanches. However, this preference was lost in early adulthood. Interestingly, in MS males but not females this preference was nonexistent already in juvenescence. There were no MS-related changes in the avalanche occurrence rate in two-region avalanches for both ages and sexes. However, in young adults, prefrontal clusters in PFC to BLA avalanches were larger in MS group and one event in the mPFC on average was followed by a smaller number of events in BLA as compared to controls indicating impaired spread of activity from the mPFC to the BLA. Importantly, the effects were specific for male animals. Spread of activity from the BLA to the mPFC was unaffected by MS in all groups.

## Discussion

High quality maternal care is essential for child’s development and early psychological deprivation is a potent stress factor, which can have long-lasting consequences (Nelson et al. 2019). Maternal separation is a long-established animal model of ELS with multifaceted effect on offsprings in terms of neurodevelopment, social and cognitive behaviors (Zhang et al. 2024). In this study, we investigated MS effect on neuronal avalanches in the developing prefrontal-amygdala circuit of juvenile and young adult rats using in vivo recordings.

The concept of neuronal avalanches, or spatio-temporal cascades of activity (Beggs and Plenz 2003), is widely exploited to study criticality in the brain networks. A hallmark of the systems near the critical state is a scale-free distribution of avalanche sizes which was successfully shown in the local cortical networks (Beggs and Plenz 2003; Gireesh and Plenz 2008; Stewart and Plenz 2008; Ma et al. 2019), as well as in largescale EEG and MEG data (Palva et al. 2013; Shriki et al. 2013; Zhigalov et al. 2015). In line with these results, we have shown previously (Kharybina et al. 2026) that size distributions of neuronal avalanches in the developing mPFC are described by truncated power law, however the same did not apply to the BLA where size distributions were characterized by a sharp peak corresponding to the maximum number of channels used for the avalanche detection indicative of the supercritical regime. The matching results were found in the current study for the animals after MS.

An important avalanche parameter characterizing the spread of activity is a branching ratio which estimates how many events are triggered by a single event and equals one for the critical systems (Kinouchi and Prado 1999). Current study shows that MS has a profound effect on the branching ratio in both BLA and mPFC in young adult rats, however no differences were found in juvenile pups suggesting that MS has long lasting consequences for the avalanche dynamics emerging later in life.

Prefrontal avalanches in the MS group were characterized by an elevated branching ratio for a large cluster of parameters used for avalanche detection, indicating intensified propagation of the activity through the system. This contradicts studies reporting ELS-related hypoactivation of the prefrontal cortex (Levine et al. 2007; Levine et al. 2008; Ishikawa et al. 2015), reduced thickness of the myelin sheath (Makinodan et al. 2012; Yang et al. 2017) and reduced dendritic length and branching (Pascual and Zamora-León 2007; Chocyk et al. 2013) in the mPFC, pointing towards impaired activity propagation. However, other studies report MS-induced increase in the dendritic length and branching (Muhammad et al. 2012) and elevated spine formation (Bock et al. 2005) in the mPFC, which is in agreement with our results. Possible explanation for these discrepancies is high specificity of the stress effects depending on the type of stress (Muhammad et al. 2012; Catale et al. 2020), its severity (Hanson et al. 2012) and exact developmental stage of stress induction (Bock et al. 2005; Makinodan et al. 2012). Moreover, ELS effects could be transient (Kraszpulski et al. 2006; Haikonen et al. 2022; Donati et al. 2026) or, as observed in the current study, manifest only at the later developmental stages, so exact timing of the examination plays crucial role in the stress outcome.

In the BLA, MS led to the elevated branching ratio for the bin sizes larger than 12 ms. Intensified propagation of the activity is in line with the reported ELS-related amygdala hyperactivity (Cohen et al. 2013; Ishikawa et al. 2015) and increased myelination of axons in the BLA (Ono et al. 2008), as well as stress-induced enlarged dendritic arborization (Vyas et al. 2002; Mitra et al. 2005). For the bin sizes less than 5 ms the opposite was true. However, given propagation delays (Madadi Asl et al. 2018) this decrease in the branching ratio is unlikely to reflect changes in the spread of activity from neuron to neuron but rather in the spontaneous activity of individual neurons or processing of the common input. One of the possible causes of such changes could be MS induced functional reorganization of the GABAergic microcircuits in the BLA (Englund et al. 2021; Haikonen et al. 2022) modulating excitability of the principal neurons.

Human and animal studies reveal sex differences in stress response from the molecular to the systems level (Bangasser and Valentino 2014). In addition, developmental trajectories of brain networks differ between males and females (Giedd et al. 1996; Lenroot et al. 2007) and the same early adverse experience occur at the different developmental stages of the brain networks depending on the sex of the animal, suggesting diversity and sex-specificity of ELS outcome (Pechtel and Pizzagalli 2011). Indeed, previously (Haikonen et al. 2022) we have linked MS to the anxiety-like behavior in male but no female rats. Accordingly, observed MS-related changes in the avalanche dynamics in both BLA and mPFC were male-specific.

Functional connectivity between amygdala and PFC is characterized by developmental switch from positive to negative coupling (Gee et al. 2013a) accelerated in children after maternal deprivation (Gee et al. 2013b). Increased negative functional connectivity between amygdala and PFC was found in infants exposed to prenatal maternal depression (Posner et al. 2016). Consistent with these results, we have shown previously (Haikonen et al. 2022) that in male rats MS facilitates development of PFC projections to GABAergic interneurons in the BLA and leads to aberrant functional prefrontal-amygdala connectivity.

Here, we asked whether these MS-induced changes in connectivity will be reflected in the spread of avalanches between two regions. In control animals, a small part of the avalanches occurred simultaneously in both regions. From them more avalanches were directed from the mPFC to the BLA than from the BLA to the mPFC in juvenile animals but not in adults (Kharybina et al. 2026). That can be related to the fact that while projections from the BLA to the mPFC emerge early in the development, their counterparts are not fully developed in juvenescence (Bouwmeester et al. 2002b; Bouwmeester et al. 2002a). Current study revealed that in MS males, but not females, the difference between the rates of the avalanches with different directionality disappears already in the juvenescence, in line with the previously reported male-specific faster maturation of the prefrontal projections (Haikonen et al. 2022). Moreover, our results show male-specific impaired avalanche transfer from the mPFC to the BLA. This can be explained by increased GABAergic drive in the BLA following prefrontal inputs in MS males (Haikonen et al. 2022) and aligns with the data on ELS-caused abnormal functional connectivity between BLA and PFC (Posner et al. 2016; Haikonen et al. 2022; Donati et al. 2026). However, while changes in GABAergic drive and functional connectivity were seen in juvenile stage, MS related changes in avalanche transfer manifested only in young adulthood. Importantly, spread of avalanches from the BLA to the mPFC was not affected by MS despite the existent evidence of the accelerated development of the amygdala projections to the prefrontal cortex following ELS (Honeycutt et al. 2020; Song et al. 2025). Yet, if these changes relate to the Glu- or GABAergic neuron targeting projections, or both, has not been established. Hence, our results on the unaffected spread of the avalanches from the BLA to the mPFC do not contradict these findings. Note, however, that we have not shown causal relationship between nLFP clusters in the regions. We cannot rule out the possibility that some of the clusters are caused by common input from another region, e.g. hippocampus which has connections to both regions (Pitkänen et al. 2000; Parent et al. 2010).

This study has one major practical limitation. Postsurgical stress (Goldkuhl et al. 2010) especially at the most vulnerable juvenile stage can easily override mild MS effects. To avoid this, in vivo multielectrode recordings were performed on lightly urethane-anaesthetized animals right after the surgery. While urethane anaesthesia resembles natural sleep (Clement et al. 2008; Mondino et al. 2022), we cannot ignore the possibility that it can have an impact on the activity spread. In vivo studies on anaesthetized animals (Gireesh and Plenz 2008; Hahn et al. 2010) showed power-law scaling of cortical avalanche size distributions, as we see it in our results as well. However, to our knowledge previously there were no studies on amygdaloid avalanches. Besides, protracted development of the mPFC – amygdala circuitry extends well into adulthood (Kolk and Rakic 2022) and beyond time investigated in the current study. Thus, it would be interesting to repeat our experiments on freely behaving adult animals providing sufficient time for post-surgery recovery and habituation and compare them to anaesthetized adult animals.

To summarize, current study was the first to demonstrate male-specific effect of ELS on neuronal avalanches occurring in the developing mPFC – BLA circuitry. Our results show MS-induced increase in the activity spread in both regions and suggest impaired spread of activity from the mPFC to the BLA. In addition, following MS activity spread between the regions showed mature-like balance at earlier developmental stages.

## Conflict of interest statement

The authors declare no competing financial interests.

## Acknowledgements

This study was financially supported by Jane and Aatos Erkko Foundation and the Research Council of Finland, grant number 361235.

## Author contribution

Z.K.: performed research, analyzed data, wrote the paper. Z.K., M.P., S.P contributed analytic tools. M.P., S.P., S.L., H.H., T.T.: designed research. H.H., T.T.: provided resources for the experimental work, conceptualized and supervised the project. S.L. and T.T.: edited the paper.

---

**Figure S1.**
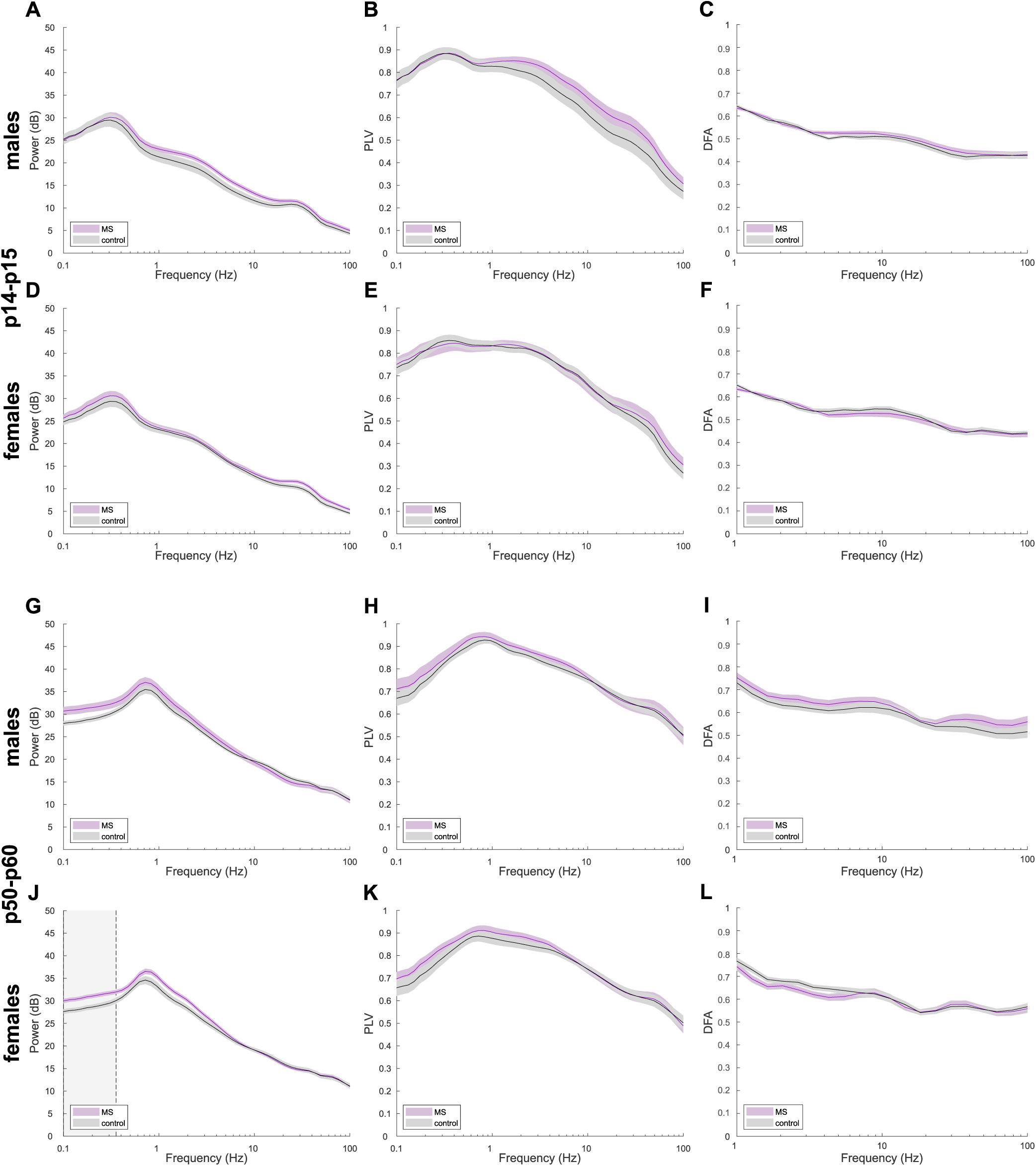
Mean power spectra (A) calculated for 16 channels in the mPFC and mean phase locking value (B) calculated between all pairs from the same 16 channels, averaged across juvenile males. (C) DFA exponents calculated for 5 nearby channels in the mPFC averaged across animals. In (A-C) shaded area represents SEM, N = 15 for both control (*black*) and MS (*magenta*) groups. (D-F) Same as (A-C) for juvenile females. N = 15 for both control and MS groups. (G-I) Same as (A-C) for adolescent males. In (G), (H) N = 17, 12, in (I) N = 21, 18 for control and MS groups, respectfully. (J-L) Same as (A-C) for adolescent females. In (J, L) N = 15, 19, in (I) N = 19, 26 for control and MS groups, respectfully. In MS adolescent females, power in infra-low band is significantly higher compared to control group (grey rectangle in (J), p < 0.05, Wilcoxon rank sum test with FDR correction, q < 0.2).

**Figure S2.**
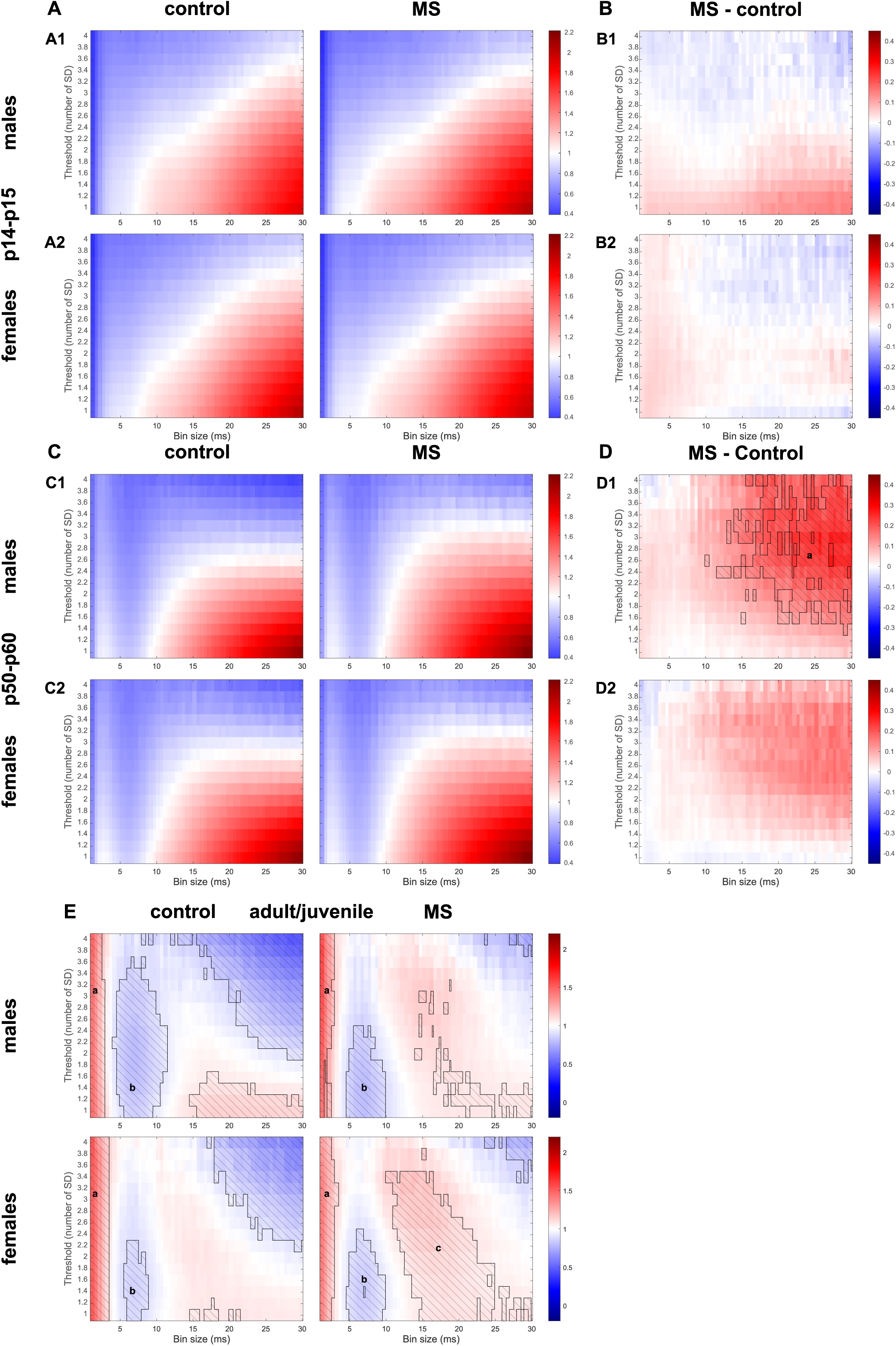
Mean branching ratio heatmaps for avalanches calculated with multiple thresholds and bin sizes in the mPFC averaged across animals. (A) Mean branching ratio calculated for juvenile males (A1) and females (A2) in control (*left*) and MS (*right*) groups. (B) Difference between mean branching ratio in MS and control groups for juvenile males (B1) and females (B2). In (A1, B1) N = 15 in both control and MS groups, in (A2, B2) N = 15 for both control and MS groups. (E, F) Same as (A, B) for young adult males (C1, D1) and females (C2, D2). In (C1, D1) N = 17, 12 for control and MS groups, respectively. Here and below hatched areas correspond to significant differences in mean branching ratio. In adolescent animals MS leads to significant increase in branching ratio for large bin sizes (significant cluster **a**: size s = 624, maximal probabilistic index *maxP(MS>control)* = 0.77; p < 0.05, Wilcoxon rank sum test with FDR correction, q < 0.2). In (C2, D2) N = 15, 19 for control and MS groups, respectively. (E) Ratio between mean branching ratio in young adult and juvenile males (E1) and females (E2) from control (*left*) and MS (*right*) groups. In males there is a significant developmental increase for small bin sizes less than 3 ms (significant cluster **a**: s = 152, *maxP(adult>juvenile)* = 0.9 and s = 125, *maxP(adult>juvenile)* = 0.89 for control and MS groups, respectively) and decrease for bin sizes between 5 and 10 ms (significant cluster **b**: s = 372, *maxP(adult<juvenile)* = 0.95 and s = 170, *maxP(adult<juvenile)* = 0.97 for control and MS groups, respectively; p < 0.05, Wilcoxon rank sum test with FDR correction, q < 0.05), developmental changes in both clusters in MS animals are less pronounced. In females there is a significant developmental increase for small bin sizes less than 3 ms (significant cluster a: s = 208, *maxP(adult>juvenile)* = 0.9 and s = 158, *maxP(adult>juvenile)* = 0.88 for control and MS groups, respectively), decrease for bin sizes between 5 and 10 ms more pronounced in MS animals (significant cluster b: s = 93, *maxP(adult<juvenile)* = 0.82 and s = 154, *maxP(adult<juvenile)* = 0.84 for control and MS groups, respectively), and increase for bin sizes between 12 and 25 ms significant only in MS animals (significant cluster c: s = 602, *maxP(adult>juvenile)* = 0.87 for MS group). For all significant clusters p < 0.05, Wilcoxon rank sum test with FDR correction, q < 0.05.

**Figure S3.**
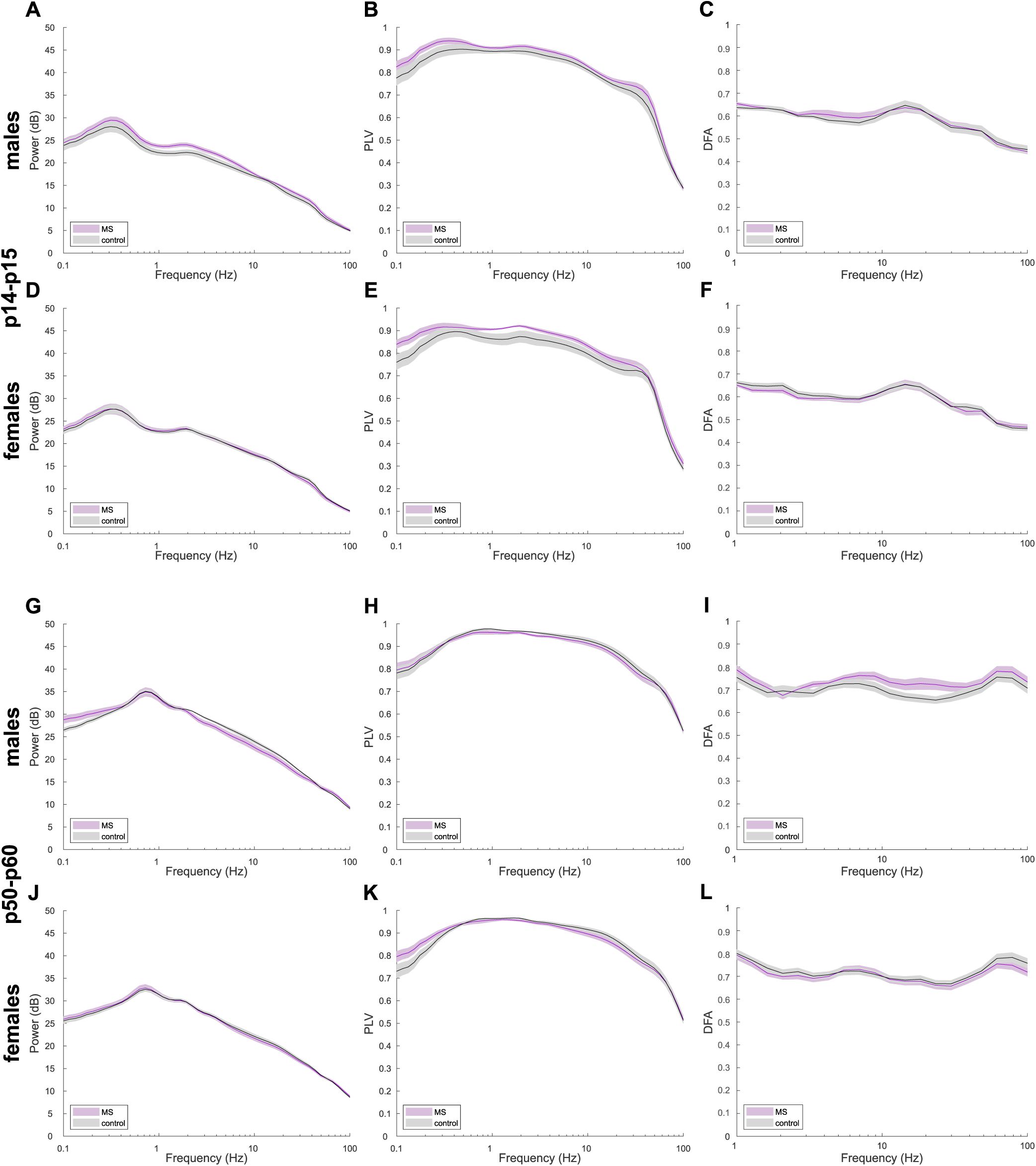
Mean power spectra (A) calculated for 16 channels in the BLA and mean phase locking value (B) calculated between all pairs from the same 16 channels, averaged across juvenile males. (C) DFA exponents calculated for 5 nearby channels in the BLA averaged across animals. In (A-C) the shaded area represents SEM. In (A, B) N = 11, 13, in C N = 15, 17 for control (*black*) and MS (*magenta*) groups, respectfully. (D-F) Same as (A-C) for juvenile females. In (D, E) N = 17, 11; in (F) N = 18, 17 for control and MS groups, respectfully. (G-I) Same as (A-C) for adolescent males. In (G, H) N = 13, 12 for control and MS groups, respectfully; in (I) N = 18 for both control and MS groups. (J-L) Same as (A-C) for adolescent females. In (J, K) N = 15, 18, in (L) N = 18, 24 for control and MS groups, respectfully.

**Figure S4.**
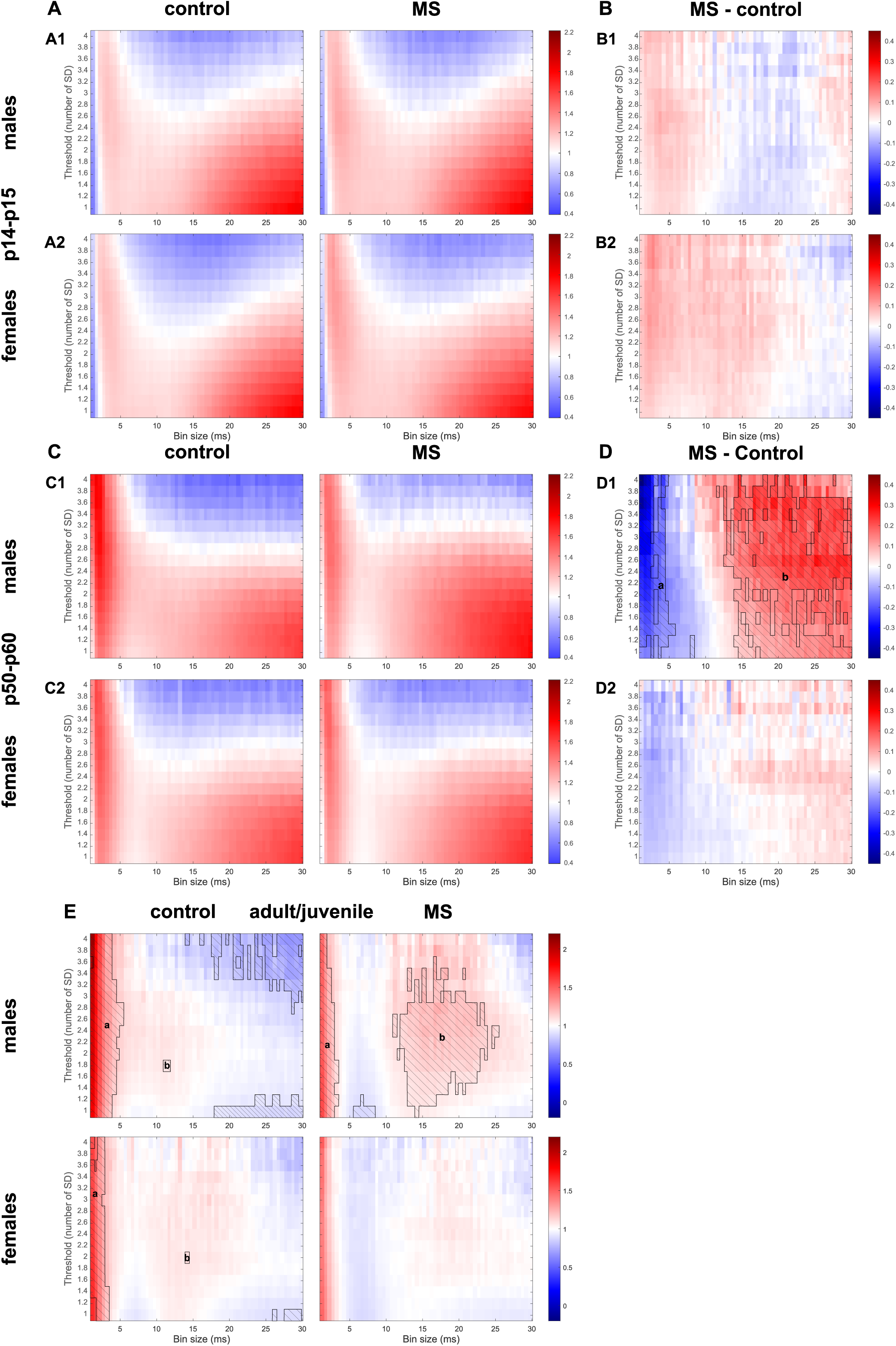
Mean branching ratio heatmaps for avalanches calculated with multiple thresholds and bin sizes in the BLA averaged across animals. (A) Mean branching ratio calculated for juvenile males (A1) and females (A2) in control (*left*) and MS (*right*) groups. (B) Difference between mean branching ratio in MS and control groups for juvenile males (B1) and females (B2). In (A1, B1) N = 11, 13 for control and MS groups, respectively; in (A2, B2) N = 17, 11 for control and MS groups, respectively. (C, D) Same as (A, B) for young adult males (C1, D1) and females (C2, D2). In (C1, D1) N = 13, 12 for control and MS groups, respectively. Here and below hatched areas correspond to significant differences in mean branching ratio. In young adult animals MS leads to significant decrease in branching ratio for small bin sizes less than 5 ms and increase for large bin sizes more than 13 ms (significant cluster **a**: size s = 245, maximal probabilistic index *maxP(MS<control)* = 0.83; significant cluster **b**: size s = 953, *maxP(MS>control)* = 0.86; p < 0.05, Wilcoxon rank sum test with FDR correction, q < 0.05). In (C2, D2) N = 15, 18 for control and MS groups, respectively. (E) Ratio between mean branching ratio in young adult and juvenile males (E1) and females (E2) from control (*left*) and MS (*right*) groups. In males there is a significant developmental increase for small bin sizes less than 5 ms less pronounced in MS group (significant cluster **a**: s = 282, *maxP(adult>juvenile)* = 0.96 and s = 148, *maxP(adult>juvenile)* = 0.88 in control and MS groups, respectively) and increase for bin sizes between 13 and 22 ms significant only in MS group (significant cluster **b**: size s = 483, *maxP(adult>juvenile)* = 0.99). In females there is a developmental increase for small bin sizes less than 3 ms significant only in control animals (significant cluster **a**: s = 149, *maxP(adult>juvenile)* = 0.86 for control group). For all significant clusters p < 0.05, Wilcoxon rank sum test with FDR correction, q < 0.05.

**Figure S5.**
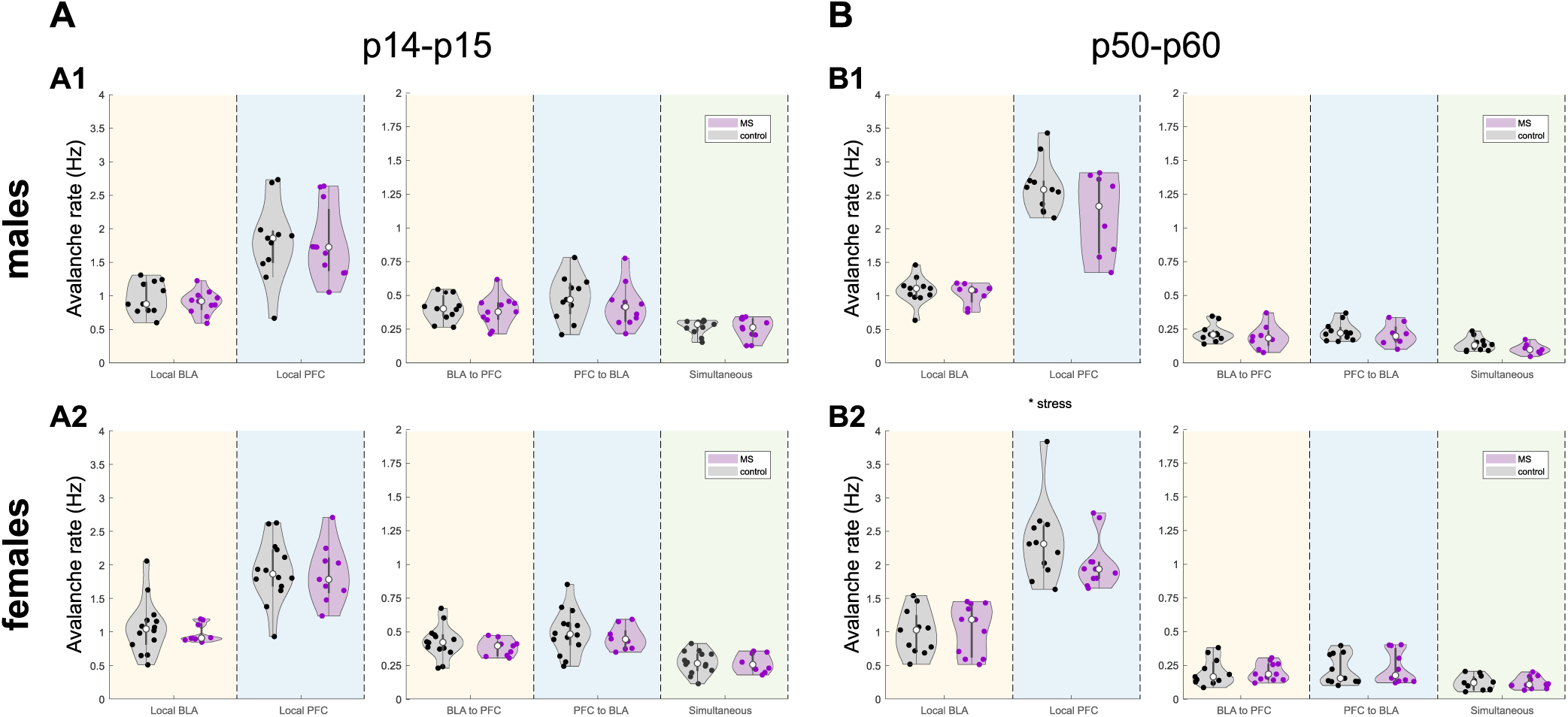
Occurrence rate of avalanches with different localization for control (*black*) and MS (*magenta*) groups in juvenile (A) and young adult (B) males (A1, B1) and females (A2, B2). (A) There are no MS related differences in all types of avalanches in juvenile animals. From left to right main effects: stress F = 0.893, 0.0173, 0.913, 1.209, 0.101, P = 0.350, 0.896, 0.345, 0.278, 0.752; sex F = 1.132, 0.354, 0.0748, 0.267, 0.149, P = 0.294, 0.555, 0.786, 0.608, 0.702; interaction F = 0.0122, 0.0158, 0.238, 0.00243, 0.00917, P = 0.912, 0.900, 0.629, 0.961, 0.924. Two-way ANOVA, in males N = 11 in both control and MS groups, in females N = 14, 9 in control and MS group, respectively. (B) MS leads to a decreased occurrence rate in local PFC avalanches and has no effect on other types of avalanches. From left to right main effects: stress F = 0.0265, 6.218, 0.287, 0.0305, 1.344, P = 0.872, 0.017, 0.595, 0.862, 0.254; sex F = 0.0778, 2.047, 0.0905, 0.0640, 0.0331, P = 0.782, 0.161, 0.765, 0.802, 0.857; interaction F = 0.195, 0.205, 0.820, 0.424, 1.321, P = 0.661, 0.653, 0.371, 0.519, 0.258. Two-way ANOVA on ranks (local BLA), raw (simultaneous avalanches) or log-transformed (all other) data, in males N = 11, 8 in control and MS group, respectively, in females N = 11 in both control and MS groups. Holm-Sidak all-pairwise comparison for local PFC avalanches: there is no stress effect in both males (t = 1.999, p = 0.053) and females (t = 1.509, p = 0.140) and no sex effect in both control (t = 1.393, p = 0.172) and MS (t = 0.664, p = 0.511) group.

**Figure S6.**
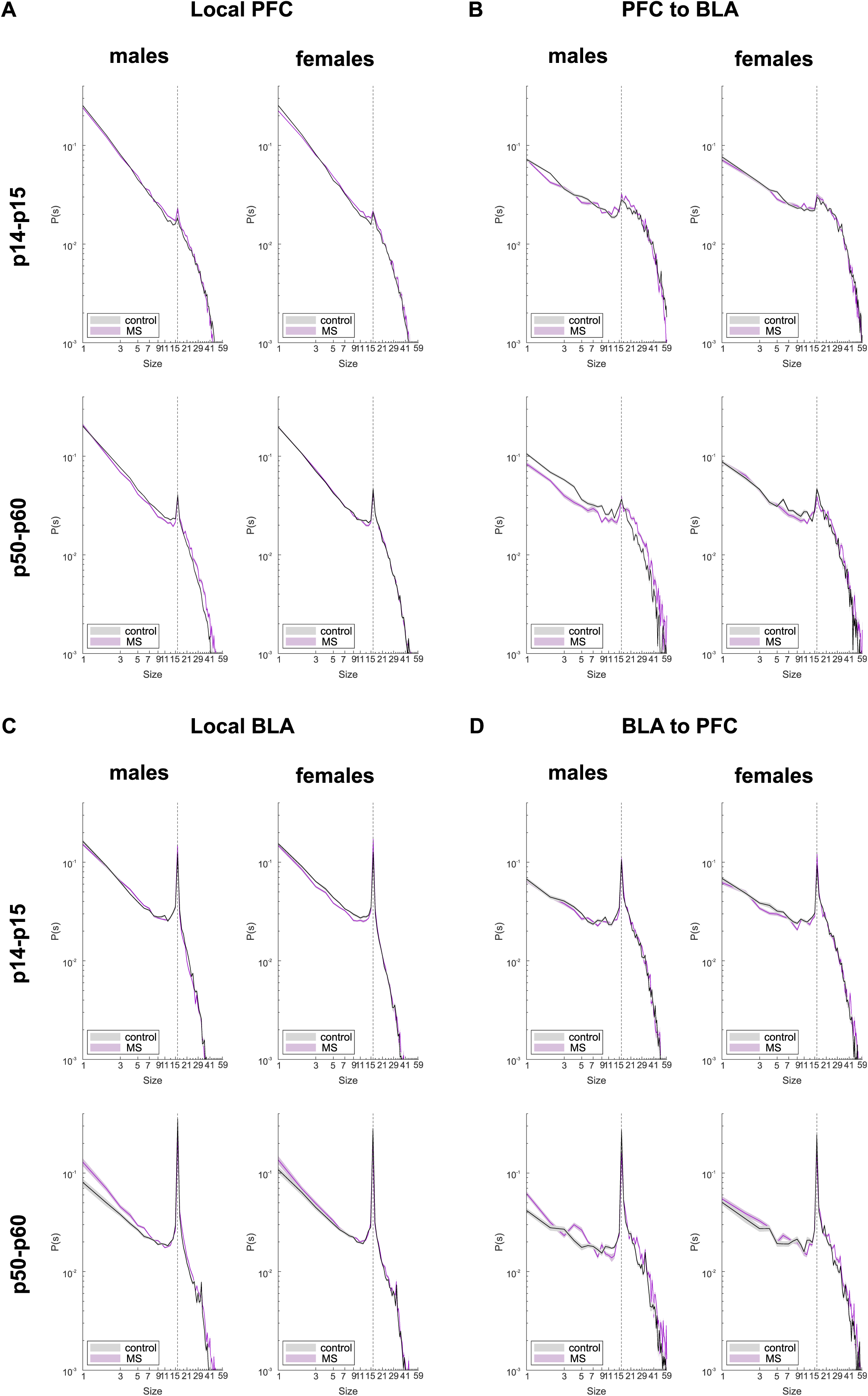
Mean size distribution of prefrontal nLFP clusters from local PFC avalanches (A), PFC to BLA avalanches (B), and amygdala nLFP clusters from local BLA avalanches (C) and BLA to PFC avalanches (D) for male (*left panel*) and female (*right panel*) juvenile (*top panel*) and young adult (*bottom panel*) animals from control (*black*) and MS (*magenta*) groups. Shaded area represents SEM.

## Notes

### Competing Interest Statement

The authors have declared no competing interest.

## References

Acion, L., Peterson, J.J., Temple, S. and Arndt, S. 2006. Probabilistic index: An intuitive non-parametric approach to measuring the size of treatment effects. Statistics in Medicine 25(4). doi: 10.1002/sim.2256.

Bak, P., Tang, C. and Wiesenfeld, K. 1987. Self-organized criticality: An explanation of the 1/f noise. Physical Review Letters 59(4). doi: 10.1103/PhysRevLett.59.381.

Bangasser, D.A. and Valentino, R.J. 2014. Sex differences in stress-related psychiatric disorders: Neurobiological perspectives. Frontiers in Neuroendocrinology 35(3). doi: 10.1016/j.yfrne.2014.03.008.

Beggs, J.M. and Plenz, D. 2003. Neuronal avalanches in neocortical circuits. The Journal of neuroscience 23(35), pp. 11167–11177. doi: 23/35/11167.

De Benedictis, A., Rossi-Espagnet, M.C., de Palma, L., Sarubbo, S. and Marras, C.E. 2023. Structural networking of the developing brain: from maturation to neurosurgical implications. Frontiers in Neuroanatomy 17. doi: 10.3389/fnana.2023.1242757.

Bock, J., Gruss, M., Becker, S. and Braun, K. 2005. Experience-induced changes of dendritic spine densities in the prefrontal and sensory cortex: Correlation with developmental time windows. Cerebral Cortex 15(6). doi: 10.1093/cercor/bhh181.

Bouwmeester, H., Smits, K. and Van Ree, J.M. 2002a. Neonatal development of projections to the basolateral amygdala from prefrontal and thalamic structures in rat. Journal of Comparative Neurology 450(3). doi: 10.1002/cne.10321.

Bouwmeester, H., Wolterink, G. and Van Ree, J.M. 2002b. Neonatal development of projections from the basolateral amygdala to prefrontal, striatal, and thalamic structures in the rat. Journal of Comparative Neurology 442(3). doi: 10.1002/cne.10084.

Catale, C., Bussone, S., Lo Iacono, L., Viscomi, M.T., Palacios, D., Troisi, A. and Carola, V. 2020. Exposure to different early-life stress experiences results in differentially altered DNA methylation in the brain and immune system. Neurobiology of Stress 13. doi: 10.1016/j.ynstr.2020.100249.

Chocyk, A., Bobula, B., Dudys, D., Przyborowska, A., Majcher-Maślanka, I., Hess, G. and Wedzony, K. 2013. Early-life stress affects the structural and functional plasticity of the medial prefrontal cortex in adolescent rats. European Journal of Neuroscience 38(1). doi: 10.1111/ejn.12208.

Clement, E.A., Richard, A., Thwaites, M., Ailon, J., Peters, S. and Dickson, C.T. 2008. Cyclic and sleep-like spontaneous alternations of brain state under urethane anaesthesia. PLoS ONE 3(4). doi: 10.1371/journal.pone.0002004.

Cocchi, L., Gollo, L.L., Zalesky, A. and Breakspear, M. 2017. Criticality in the brain: A synthesis of neurobiology, models and cognition. Progress in Neurobiology 158, pp. 132–152. doi: 10.1016/j.pneurobio.2017.07.002.

Cohen, M.M., Jing, D., Yang, R.R., Tottenham, N., Lee, F.S. and Casey, B.J. 2013. Early-life stress has persistent effects on amygdala function and development in mice and humans. Proceedings of the National Academy of Sciences of the United States of America 110(45). doi: 10.1073/pnas.1310163110.

Dennis, E.L. and Thompson, P.M. 2013. Mapping connectivity in the developing brain. International Journal of Developmental Neuroscience 31(7). doi: 10.1016/j.ijdevneu.2013.05.007.

Donati, A., Vedele, F. and Hartung, H. 2026. Early-life stress impairs development of functional interactions and neuronal activity within prefrontal-amygdala networks *in vivo*. Molecular Psychiatry. doi: 10.1038/s41380-026-03448-z.

Edmiston, E.E., Wang, F., Mazure, C.M., Guiney, J., Sinha, R., Mayes, L.C. and Blumberg, H.P. 2011. Corticostriatal-limbic gray matter morphology in adolescents with self-reported exposure to childhood maltreatment. Archives of Pediatrics and Adolescent Medicine 165(12). doi: 10.1001/archpediatrics.2011.565.

Englund, J. et al. 2021. Downregulation of kainate receptors regulating GABAergic transmission in amygdala after early life stress is associated with anxiety-like behavior in rodents. Translational Psychiatry 11(1). doi: 10.1038/s41398-021-01654-7.

Famularo, R., Kinscherff, R. and Fenton, T. 1992. Psychiatric Diagnoses of Maltreated Children: Preliminary Findings. Journal of the American Academy of Child and Adolescent Psychiatry 31(5). doi: 10.1097/00004583-199209000-00013.

Gee, D.G. et al. 2013a. A developmental shift from positive to negative connectivity in human amygdala-prefrontal circuitry. Journal of Neuroscience 33(10). doi: 10.1523/JNEUROSCI.3446-12.2013.

Gee, D.G. et al. 2013b. Early developmental emergence of human amygdala-prefrontal connectivity after maternal deprivation. Proceedings of the National Academy of Sciences of the United States of America 110(39). doi: 10.1073/pnas.1307893110.

Giedd, J.N. et al. 1996. Quantitative MRI of the temporal lobe, amygdala, and hippocampus in normal human development: Ages 4-18 years. Journal of Comparative Neurology 366(2). doi: 10.1002/(SICI)1096-9861(19960304)366:2<223::AID-CNE3>3.0.CO;2-7.

Gireesh, E.D. and Plenz, D. 2008. Neuronal avalanches organize as nested theta- and beta/gamma-oscillations during development of cortical layer 2/3. Proceedings of the National Academy of Sciences of the United States of America 105(21), pp. 7576–7581. doi: 10.1073/pnas.0800537105.

Goldkuhl, R., Klockars, A., Carlsson, H.E., Hau, J. and Abelson, K.S.P. 2010. Impact of surgical severity and analgesic treatment on plasma corticosterone in rats during surgery. European Surgical Research 44(2). doi: 10.1159/000264962.

Hahn, G., Petermann, T., Havenith, M.N., Yu, S., Singer, W., Plenz, D. and Nikolić, D. 2010. Neuronal avalanches in spontaneous activity in vivo. Journal of Neurophysiology 104(6). doi: 10.1152/jn.00953.2009.

Haikonen, J. et al. 2022. Aberrant cortical projections to amygdala GABAergic neurons contribute to developmental circuit dysfunction following early life stress. iScience 26(1). doi: 10.1016/j.isci.2022.105724.

Hanson, J.L. et al. 2012. Structural variations in prefrontal cortex mediate the relationship between early childhood stress and spatial working memory. Journal of Neuroscience 32(23). doi: 10.1523/JNEUROSCI.0307-12.2012.

Hanson, J.L. et al. 2015. Behavioral problems after early life stress: Contributions of the hippocampus and amygdala. Biological Psychiatry 77(4). doi: 10.1016/j.biopsych.2014.04.020.

Hardstone, R., Poil, S.S., Schiavone, G., Jansen, R., Nikulin, V. V, Mansvelder, H.D. and Linkenkaer-Hansen, K. 2012. Detrended fluctuation analysis: a scale-free view on neuronal oscillations. Frontiers in physiology 3, p. 450. doi: 10.3389/fphys.2012.00450.

Van Harmelen, A.L. et al. 2010. Reduced medial prefrontal cortex volume in adults reporting childhood emotional maltreatment. Biological Psychiatry 68(9). doi: 10.1016/j.biopsych.2010.06.011.

Van Harmelen, A.L. et al. 2013. Enhanced amygdala reactivity to emotional faces in adults reporting childhood emotional maltreatment. Social Cognitive and Affective Neuroscience 8(4). doi: 10.1093/scan/nss007.

Van Harmelen, A.L. et al. 2014. Hypoactive medial prefrontal cortex functioning in adults reporting childhood emotional maltreatment. Social Cognitive and Affective Neuroscience 9(12). doi: 10.1093/scan/nsu008.

Heiney, K., Huse Ramstad, O., Fiskum, V., Christiansen, N., Sandvig, A., Nichele, S. and Sandvig, I. 2021. Criticality, Connectivity, and Neural Disorder: A Multifaceted Approach to Neural Computation. Frontiers in Computational Neuroscience 15. doi: 10.3389/fncom.2021.611183.

Honeycutt, J.A. et al. 2020. Altered corticolimbic connectivity reveals sex-specific adolescent outcomes in a rat model of early life adversity. eLife 9. doi: 10.7554/eLife.52651.

Ishikawa, J., Nishimura, R. and Ishikawa, A. 2015. Early-life stress induces anxiety-like behaviors and activity imbalances in the medial prefrontal cortex and amygdala in adult rats. European Journal of Neuroscience 41(4). doi: 10.1111/ejn.12825.

Kharybina, Z., Palva, M., Palva, S., Lauri, S., Hartung, H. and Taira, T. 2026. Different developmental trajectories of critical dynamics in prefrontal cortex–amygdala circuitry. bioRxiv. doi: 10.64898/2026.02.13.705704.

Kinouchi, O. and Prado, C.P.C. 1999. Robustness of scale invariance in models with self-organized criticality. *Physical Review E - Statistical Physics, Plasmas*, Fluids, and Related Interdisciplinary Topics 59(5). doi: 10.1103/PhysRevE.59.4964.

Kolk, S.M. and Rakic, P. 2022. Development of prefrontal cortex. Neuropsychopharmacology 47(1), pp. 41–57. doi: 10.1038/s41386-021-01137-9.

Kraszpulski, M., Dickerson, P.A. and Salm, A.K. 2006. Prenatal stress affects the developmental trajectory of the rat amygdala. Stress 9(2). doi: 10.1080/10253890600798109.

Lenroot, R.K. et al. 2007. Sexual dimorphism of brain developmental trajectories during childhood and adolescence. NeuroImage 36(4). doi: 10.1016/j.neuroimage.2007.03.053.

Levine, J.B. et al. 2008. Isolation rearing impairs wound healing and is associated with increased locomotion and decreased immediate early gene expression in the medial prefrontal cortex of juvenile rats. Neuroscience 151(2). doi: 10.1016/j.neuroscience.2007.10.014.

Levine, J.B., Youngs, R.M., MacDonald, M.L., Chu, M., Leeder, A.D., Berthiaume, F. and Konradi, C. 2007. Isolation rearing and hyperlocomotion are associated with reduced immediate early gene expression levels in the medial prefrontal cortex. Neuroscience 145(1). doi: 10.1016/j.neuroscience.2006.11.063.

Lupien, S.J., McEwen, B.S., Gunnar, M.R. and Heim, C. 2009. Effects of stress throughout the lifespan on the brain, behaviour and cognition. Nature Reviews Neuroscience 10(6). doi: 10.1038/nrn2639.

Ma, Z., Turrigiano, G.G., Wessel, R. and Hengen, K.B. 2019. Cortical Circuit Dynamics Are Homeostatically Tuned to Criticality In Vivo. Neuron 104(4), pp. 655–664.e4. doi: S0896-6273(19)30737-8.

Madadi Asl, M., Valizadeh, A. and Tass, P.A. 2018. Dendritic and Axonal Propagation Delays May Shape Neuronal Networks With Plastic Synapses. Frontiers in Physiology 9. doi: 10.3389/fphys.2018.01849.

Makinodan, M., Rosen, K.M., Ito, S. and Corfas, G. 2012. A critical period for social experience-dependent oligodendrocyte maturation and myelination. Science 337(6100). doi: 10.1126/science.1220845.

Makris, G., Eleftheriades, A. and Pervanidou, P. 2023. Early Life Stress, Hormones, and Neurodevelopmental Disorders. Hormone Research in Paediatrics 96(1). doi: 10.1159/000523942.

McCrory, E., De Brito, S.A. and Viding, E. 2012. The link between child abuse and psychopathology: A review of neurobiological and genetic research. Journal of the Royal Society of Medicine 105(4). doi: 10.1258/jrsm.2011.110222.

McCrory, E.J., De Brito, S.A., Sebastian, C.L., Mechelli, A., Bird, G., Kelly, P.A. and Viding, E. 2011. Heightened neural reactivity to threat in child victims of family violence. Current Biology 21(23). doi: 10.1016/j.cub.2011.10.015.

McGinty, V.B. and Grace, A.A. 2008. Selective activation of medial prefrontal-to-accumbens projection neurons by amygdala stimulation and pavlovian conditioned stimuli. Cerebral Cortex 18(8). doi: 10.1093/cercor/bhm223.

McLaughlin, K.A., Sheridan, M.A., Winter, W., Fox, N.A., Zeanah, C.H. and Nelson, C.A. 2014. Widespread reductions in cortical thickness following severe early-life deprivation: A neurodevelopmental pathway to attention-deficit/hyperactivity disorder. Biological Psychiatry 76(8). doi: 10.1016/j.biopsych.2013.08.016.

Mehta, M.A. et al. 2009. Amygdala, hippocampal and corpus callosum size following severe early institutional deprivation: the English and Romanian Adoptees study pilot. Journal of child psychology and psychiatry, and allied disciplines 50(8). doi: 10.1111/j.1469-7610.2009.02084.x.

Mitra, R., Jadhav, S., McEwen, B.S., Vyas, A. and Chattarji, S. 2005. Stress duration modulates the spatiotemporal patterns of spine formation in the basolateral amygdala. Proceedings of the National Academy of Sciences of the United States of America 102(26). doi: 10.1073/pnas.0504011102.

Mondino, A. et al. 2022. Urethane anaesthesia exhibits neurophysiological correlates of unconsciousness and is distinct from sleep. European Journal of Neuroscience 59(4), pp. 483–501. doi: 10.1111/ejn.15690.

Muhammad, A., Carroll, C. and Kolb, B. 2012. Stress during development alters dendritic morphology in the nucleus accumbens and prefrontal cortex. Neuroscience 216. doi: 10.1016/j.neuroscience.2012.04.041.

Nelson, C.A., Zeanah, C.H. and Fox, N.A. 2019. How early experience shapes human development: The case of psychosocial deprivation. Neural Plasticity 2019. doi: 10.1155/2019/1676285.

Nieves, G.M., Bravo, M., Baskoylu, S. and Bath, K.G. 2020. Early life adversity decreases pre-adolescent fear expression by accelerating amygdala PV cell development. eLife 9. doi: 10.7554/eLife.55263.

Ono, M., Kikusui, T., Sasaki, N., Ichikawa, M., Mori, Y. and Murakami-Murofushi, K. 2008. Early weaning induces anxiety and precocious myelination in the anterior part of the basolateral amygdala of male Balb/c mice. Neuroscience 156(4). doi: 10.1016/j.neuroscience.2008.07.078.

Palva, J.M., Zhigalov, A., Hirvonen, J., Korhonen, O., Linkenkaer-Hansen, K. and Palva, S. 2013. Neuronal long-range temporal correlations and avalanche dynamics are correlated with behavioral scaling laws. Proceedings of the National Academy of Sciences of the United States of America 110(9), pp. 3585–3590. doi: 10.1073/pnas.1216855110.

Parent, M.A., Wang, L., Su, J., Netoff, T. and Yuan, L.L. 2010. Identification of the hippocampal input to medial prefrontal cortex in vitro. Cerebral Cortex 20(2). doi: 10.1093/cercor/bhp108.

Pascual, R. and Zamora-León, S.P. 2007. Effects of neonatal maternal deprivation and postweaning environmental complexity on dendritic morphology of prefrontal pyramidal neurons in the rat. Acta Neurobiologiae Experimentalis 67(4). doi: 10.55782/ane-2007-1663.

Pechtel, P. and Pizzagalli, D.A. 2011. Effects of early life stress on cognitive and affective function: An integrated review of human literature. Psychopharmacology 214(1). doi: 10.1007/s00213-010-2009-2.

Pirkola, S. et al. 2005. Childhood adversities as risk factors for adult mental disorders. Results from the Health 2000 study. Social Psychiatry and Psychiatric Epidemiology 40(10). doi: 10.1007/s00127-005-0950-x.

Pitkänen, A., Pikkarainen, M., Nurminen, N. and Ylinen, A. 2000. Reciprocal connections between the amygdala and the hippocampal formation, perirhinal cortex, and postrhinal cortex in rat. In: Annals of the New York Academy of Sciences. doi: 10.1111/j.1749-6632.2000.tb06738.x.

Posner, J. et al. 2016. Alterations in amygdala-prefrontal circuits in infants exposed to prenatal maternal depression. Translational Psychiatry 6(11). doi: 10.1038/tp.2016.146.

Rincel, M. and Darnaudéry, M. 2020. Maternal separation in rodents: A journey from gut to brain and nutritional perspectives. Proceedings of the Nutrition Society 79(1). doi: 10.1017/S0029665119000958.

Shriki, O. et al. 2013. Neuronal avalanches in the resting MEG of the human brain. Journal of Neuroscience 33(16). doi: 10.1523/JNEUROSCI.4286-12.2013.

Song, J., Younus, M., Long, H., Walker, C.D. and Wong, T.P. 2025. Layer-Specific Glutamatergic Inputs and Parvalbumin Interneurons Modulate Early Life Stress-Induced Alterations in Prefrontal Glutamate Release during Fear Conditioning in Pre-adolescent Rats. eNeuro 12(11). doi: 10.1523/ENEURO.0073-25.2025.

Stewart, C. V. and Plenz, D. 2008. Homeostasis of neuronal avalanches during postnatal cortex development in vitro. Journal of Neuroscience Methods 169(2). doi: 10.1016/j.jneumeth.2007.10.021.

Storey, J.D. 2002. A direct approach to false discovery rates. Journal of the Royal Statistical Society. Series B: Statistical Methodology 64(3). doi: 10.1111/1467-9868.00346.

Tau, G.Z. and Peterson, B.S. 2010. Normal development of brain circuits. Neuropsychopharmacology 35(1). doi: 10.1038/npp.2009.115.

Tottenham, N. et al. 2010. Prolonged institutional rearing is associated with atypically large amygdala volume and difficulties in emotion regulation. Developmental Science 13(1). doi: 10.1111/j.1467-7687.2009.00852.x.

Tottenham, N. 2020. Early Adversity and the Neotenous Human Brain. Biological Psychiatry 87(4). doi: 10.1016/j.biopsych.2019.06.018.

Tottenham, N., Hare, T.A., Millner, A., Gilhooly, T., Zevin, J.D. and Casey, B.J. 2011. Elevated amygdala response to faces following early deprivation. Developmental Science 14(2). doi: 10.1111/j.1467-7687.2010.00971.x.

Vyas, A., Mitra, R., Shankaranarayana Rao, B.S. and Chattarji, S. 2002. Chronic stress induces contrasting patterns of dendritic remodeling in hippocampal and amygdaloid neurons. Journal of Neuroscience 22(15). doi: 10.1523/jneurosci.22-15-06810.2002.

Yamamoto, T. et al. 2017. Increased amygdala reactivity following early life stress: A potential resilience enhancer role. BMC Psychiatry 17(1). doi: 10.1186/s12888-017-1201-x.

Yang, Y. et al. 2017. Neonatal Maternal Separation Impairs Prefrontal Cortical Myelination and Cognitive Functions in Rats Through Activation of Wnt Signaling. Cerebral Cortex 27(5). doi: 10.1093/cercor/bhw121.

Zhang, Y., Wang, S. and Hei, M. 2024. Maternal separation as early-life stress: Mechanisms of neuropsychiatric disorders and inspiration for neonatal care. Brain Research Bulletin 217. doi: 10.1016/j.brainresbull.2024.111058.

Zhigalov, A., Arnulfo, G., Nobili, L., Palva, S. and Palva, J.M. 2015. Relationship of Fast- and Slow-Timescale Neuronal Dynamics in Human MEG and SEEG. The Journal of Neuroscience 35(13), pp. 5385–5396. doi: 10.1523/JNEUROSCI.4880-14.2015.

